# Investigation of Neurons and Microglia from *APOE3* Induced Pluripotent Stem Cells using Data-Independent Acquisitions on a ZenoTOF 7600

**DOI:** 10.1101/2025.11.02.686128

**Authors:** Jordan B. Burton, Tyne McHugh, Charles A. Schurman, Joanna Bons, Maria Sanchez, Lisa M. Ellerby, Birgit Schilling

**Affiliations:** Buck Institute for Research on Aging, Novato, CA, USA; Leonard Davis School of Gerontology, University of Southern California, Los Angeles, CA 90893, United States

## Abstract

Induced pluripotent stem cell (iPSC)-derived neurons and microglia are valuable human models for studying neurodegenerative diseases. Specifically, the apolipoprotein E4 (*APOE4*) gene is a major genetic risk factor for late-onset Alzheimer’s disease. Apolipoprotein E (*APOE*) alleles E2, E3 and E4 can be beneficial, neutral, or increase the risk of Alzheimer’s disease (AD). Here, we developed a proteomic workflow using data-independent acquisitions to provide a quantitative mass spectrometric proteome analysis, and proteomic screening assays for brain-specific cell types derived from iPSC. Protein groups were quantified in *APOE3* neurons and microglia, respectively, with ∼80% overlap. Cell type-specific markers and enriched pathways reflected the specialized functions of each cell type, such as synaptic signaling in neurons and immune and inflammatory responses in microglia. The neuron-specific markers included proteins APP, CALB1, CALB2, DLGs, GAP43, NEFL, MAPs; while microglial markers included proteins AIF1, CDs, MMP9, and ITGAM. Ultimately, the combination of robust iPSC differentiation and sensitive proteomic screening assays described here provides a valuable platform for probing the cellular mechanisms underlying neurological disorders.

**Significance:** The quantification of dysregulated proteins and pathways in patient-derived neurons and microglia can provide insights into disease etiology and progression. More broadly, this DIA approach enables deep proteome profiling of unique iPSC-derived cell models, increasing their utility for investigating disease biology and therapeutic development. We focused on iPSC models from two important cell types of the brain, excitatory neurons and microglia. We integrated the proteomes of these two cell types. These tools provide robust biological and mass spectrometric screening tools for future therapeutic interventions using disease-relevant human brain cell types or brain organoid models.

**Graphical abstract:** 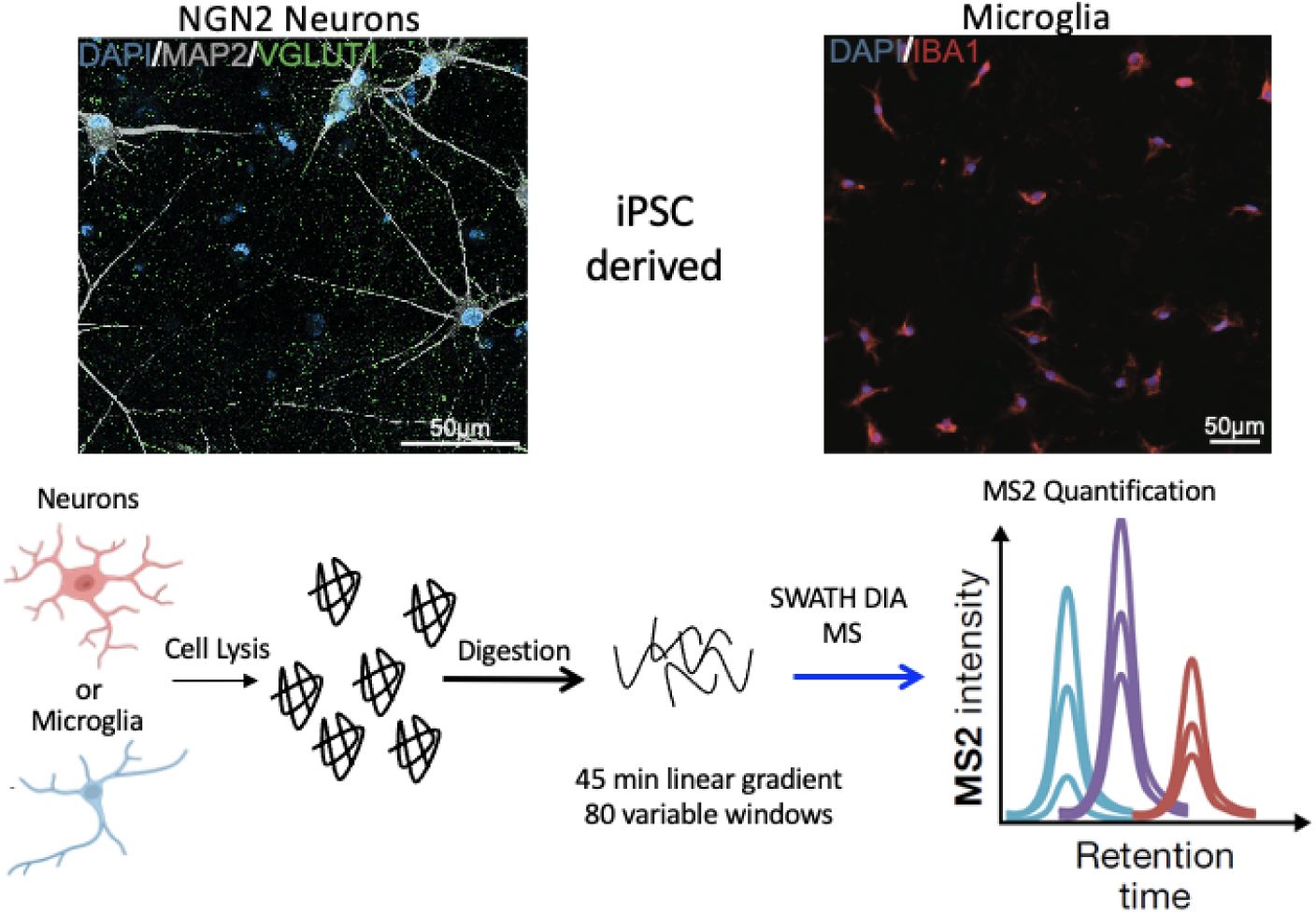

**Highlights:**

- Presentation of a proteomic workflow using data-independent acquisitions to monitor and screen proteomes of iPSC-derived brain cell types.
- Quick MS Assays to determine protein profiles of iPSC-derived neurons and microglia.
- Characterization of different cell type proteomes from *APOE3* iPSCs.
- Revealing of neuron-specific markers and microglia-specific markers by mass spectrometry.
- Step-by-step instructions for the set-up of the DIA-MS assays

## Introduction

Induced pluripotent stem cell (iPSC)-derived models have great potential for uncovering mechanisms of neurodegenerative disease pathogenesis (1). However, iPSC-based modeling has focused a wide diversity of differentiation protocols tailored to differentiation into neuronal subtypes, while protocols for generating iPSC-derived microglia are still emerging (2-9). Microglia are the primary immune cells in the central nervous system and play a role in maintaining brain homeostasis, and can contribute to neurodegenerative diseases (10). Fully characterized microglia models that recapitulate the functional properties of primary human microglia will provide important tools for investigating the microglia-specific mechanisms of neurodegeneration. However, the generation of human subtype microglia-like cells from iPSCs has faced challenges due to technical barriers. Traditional protocols have relied on either the overexpression of specific transcription factors using lentiviral gene or differentiation using small molecules, recombinant proteins, and growth factors, which demand extensive technical skills (11-13). Limitations of the latter methods have resulted in variability in microglia cell yield and limited scalability. In our study, we compared the proteome of iPSC-derived neurons and iPSC-derived microglia differentiated into their respective cell types using well established protocols (see references in methods add here). Combining iPSC technology in the context of neurodegenerative diseases with molecular technologies (proteomics/RNASeq/epigenetics) has proven to be powerful tools, as demonstrated by the Hao group (14), the NeuroLINCS consortium (15) and others (16) for ALS research.

Recent advances in mass spectrometry including both, improvements for instrumentation and data processing / machine learning, have enabled deep mining of proteomes, increased sensitivity and high quantification accuracy. Briefly, in 2012, Gillet et al. introduced label-free SWATH MS quantification (17), also referred to as data-independent acquisition (DIA-MS), opening powerful label-free acquisition solutions (18-20). Using DIA-MS researchers demonstrated the ability to quantify large numbers of proteins with quantification at the MS2 fragment ion level (high selectivity and specificity) with basically no missing values to achieve the quantitative reproducibility and specificity of targeted MS. Kitata *et al*. described a comprehensive overview early into the development of DIA-MS (21). Over the last decade, major advancements were further implemented featuring many novel software improvements, and computational innovations (22). Many of the new DIA-MS workflows have been applied for preclinical studies and for clinical investigations (23).

In this study, we employed a hybrid quadrupole-time-of-flight (QqTOF) ZenoTOF 7600 mass spectrometer (SCIEX) for DIA SWATH MS acquisitions using specific innovations. In order to maximize duty cycle efficiency and to achieve higher sensitivity, the QqTOF mass spectrometer integrates a novel trap-and-release technology (24). The linear trapping device is located between the Q2 quadrupole collision cell and the TOF accelerator. This efficiently minimizes any duty cycle ion loss when ions are released into the TOF accelerator. Ions are captured and stored in the trap after exiting the Q2 collision cell. Then, in synchronization with the TOF accelerator, the ions are released based on potential energy from higher m/z ions to lower m/z ions, such that all ions arrive at the center of the accelerator plate at the same time for efficient delivery into the TOF region for detection (24, 25). This technology improves the ion duty cycle up to >90% through this region of the MS system and enhances detection sensitivity. Recent work by the Ralser group and our grous demonstrated that the ZenoTOF 7600 enabled quantitative studies of low-level starting material (26), as well as high-throughput sample acquisitions (27, 28).

The work presented here provides a step-by-step protocol to set up the DIA SWATH MS assays on a ZenoTOF 7600 platform and subsequently analyze the data with Spectronaut (Biognosys) (see **Supplemental Protocol**). We optimized the DIA SWATH MS assay for monitoring human protein profiles from iPSC-derived neurons and microglia in the context of a human model system, featuring an apolipoprotein E3 (APOE3) allele, the most common APOE allele which typically does not pose an AD risk factor. The iPSC-derived neurons exhibited neuronal expression profiles and electrical activity indicative of functional maturation (29). The iPSC-derived microglia exhibited expression profiles and functional characteristics, including phagocytosis, cytokine secretion, and inflammatory signaling, analogous to primary microglia (12). The validated microglia model enables studies of microglia-related phenotypes and contributions in neurodegenerative diseases using an entirely human cellular system. These new integrated proteomic assays provide a tool set to biologists to employ large-scale quantitative proteomics studies with greater depth of coverage, even with limited sample amounts.

## Materials and Methods

### Reagents

Acetonitrile and water for HPLC were obtained from Burdick & Jackson (Muskegon, MI). Sodium dodecyl sulfate, triethylammonium bicarbonate buffer (TEAB), iodoacetamide, dithiothreitol (DTT), and formic acid for protein chemistry were purchased from Sigma Aldrich (St. Louis, MO). Promega (Madison WI) provided sequencing grade trypsin. S-Trap mini columns were obtained from Protifi (Fairport, NY). HLB Oasis SPE cartridges were purchased from Waters (Milford, MA). Biognosys (Schlieren, Switzerland) supplied the iRT peptides. HeLa digests were obtained from Thermo Fisher Scientific (Waltham, MA).

### NGN2 Neuron Generation

Human iPSC-derived neurons were differentiated as previously described (Wang et al., 2017) via integration of a doxycycline-inducible Neurogenin 2 (Ngn2) transgene into APOE3 carrying human iPSCs. Briefly, iPSCs were incubated with doxycycline (2 μg/mL; Sigma) for 3 days at a density of 1.5 × 10^6^ cells/well in six-well plates coated with Matrigel in knockout Dulbecco’s modified Eagle’s medium (KO-DMEM)/F12 medium containing N2 supplement, non-essential amino acids (NEAA), brain-derived neurotrophic factor (BDNF, 10 ng/mL; StemCell Technologies), neurotrophin-3 (NT3,10 ng/mL; StemCell Technologies), and Rock inhibitor (StemCell Technologies, Y27632). The medium was changed daily, and Rock inhibitor was removed from day 2. Pre-differentiated iPSCs were seeded on glass coverslips coated with poly-D-lysine (20 µg/ml; Sigma, P6407) and laminin (0.25 µg/ml; Sigma, L2020) in 24-well plates at a density of 150,000 cells per well in Neurobasal-A medium containing doxycycline (1 μg/mL), 50x B27 supplement (ThermoFisher), 400x GlutaMax (ThermoFisher), BDNF (10 ng/mL), and NT3 (10 ng/mL). Half of the medium was replaced on day 4 without doxycycline and again on day 11, and the medium volume was doubled on day 18. Thereafter, one-third of the medium was replaced weekly until the cells were harvested for proteomic analysis at day 28.

### Microglia Generation

Microglia were generated from human induced pluripotent stem cells (iPSC) as previously described (Douvaras, 2017). Briefly, iPSCs were dissociated with accutase (Gibco, A11105-01) and plated as single cells on matrigel coated plates at 15×10^3^ cells/cm2 in TeSR-E8 (StemCell Technologies, 05990) plus 10uM Rock inhibitor (StemCell Technologies, Y27632). When colonies were visible (Day 0) cells were induced to differentiation by supplementing TeSR-E8 with 80ng/ml BMP4 (R&D systems, 314-BP-010). Media supplemented with BMP4 was changed daily for 4 days. Cells were then induced with StemPro-34 SFM containing 2mM GutaMAX (Gibco, 35050-061) and supplemented with 25ng/ml bFGF (Peprotech, 100-18B), 100ng/ml SCF (R&D systems, 255-SC-010) and 80ng/ml VEGF(165) (Peprotech, 100-20). Two days later, the medium was switched to StemPro-34 containing 50ng/ml SCF, 50ng/ml IL-3 (R&D systems, 203-IL-010), 5ng/ml TPO (Peprotech, 300-18), 50ng/ml M-CSF (Peprotech, 300-25) and 50ng/ml Flt3 ligand (Peprotech, 300-19). From day 14 onward StemPro-34 media was supplemented with 50ng/ml M-CSF, 50ng/ml Flt3 ligand and 25ng/ml GM-CSF (Peprotech, 300-03). Between days 30-90 free floating cells were collected and plated in matrigel coated dishes in Advanced RPMI (Gibco,12633012) with 2 mM GlutaMAX 10ng/mL GM-CSF, and 100 ng/mL IL-34(Peprotech, 200-34). Medium was replenished every 3–4 days for at least 2 weeks before harvesting for proteomic analysis.

### Immunocytochemisty

Cells were fixed for 15 minutes with 4% paraformaldehyde in PBS and then washed 3 times with PBS followed by 1 hr incubation at room temperature in blocking buffer (0.1% Triton-X-100, 2% normal donkey serum in PBS). The cells were then incubated in blocking solution containing primary antibody for 1 hr, followed by washing and incubation with secondary fluorescent-labeled antibodies for 1 hr. Antibodies used for immunocytochemistry were those against MAP2 (NB300-213; Novus, 1:400), Vglut1 (MAB5502; Millipore, 1:1000), and Iba1 (019-19741; Fujifilm Chemicals, 1:200). Coverslips were mounted onto glass slides with Vectashield. Confocal images of the neurons were taken using a Zeiss LSM980 63x immersion oil objective, while images of microglia were taken on the Biotek.

### Proteolytic Digestion

The cell lysates were resuspended in 100 µL of 4% SDS in 50 mM triethylammonium bicarbonate buffer (TEAB) through vigorous mixing and incubated at 90 °C for 10 minutes. Samples were treated with 20 mM dithiothreitol in 50 mM TEAB at a pH of 7 at 56 °C for 10 minutes. The samples were then cooled to room temperature and allowed to sit for an additional 10 minutes. Next, the samples were alkylated with 40 mM iodacetamide in 50 mM TEAB (pH 7) at room temperature for 30 minutes in the dark. The samples were then acidified by adding 12% phosphoric acid, and a fine precipitate was formed after the addition of S-Trap buffer and mixing by inversion. The entire mixture was added to S-Trap spin columns (ProtiFi) and centrifuged at 4,000 g for 20 seconds or until fully eluted. The samples were then washed with an additional volume of S-Trap buffer. Next, the samples were incubated with sequencing grade trypsin (Promega) dissolved in 50 mM TEAB (pH 7) at a 1:25 enzyme:protein ratio for 1 hour at 47 °C. Additional trypsin was added at the same ratio and the proteins were digested at 37 °C overnight. The peptides were eluted from the S-Trap columns and dried by centrifugal evaporation, and then resuspended in 1% formic acid in water and desalted using 10 mg Oasis SPE cartridges (Waters). The desalted elutions were dried by centrifugal evaporation and resuspended in 0.2% formic acid in water. Finally, indexed Retention Time Standards (iRT, Biognosys) were added to each sample according to the manufacturer’s instructions (30).

### Chromatographic separation and subsequent mass spectrometric analysis

Reverse-phase HPLC-MS/MS data was collected using a Waters M-Class HPLC (Waters, Massachusetts, MA) connected to a ZenoTOF 7600 (SCIEX, Redwood City, CA) with an OptiFlow Turbo V Ion Source (SCIEX). A microelectrode with a flow rate of 1-10 µL/min was used. The solvent system consisted of 0.1% formic acid in water (solvent A) and 99.9% acetonitrile, 0.1% formic acid in water (solvent B). Digested peptides were loaded on a Luna Micro C18 trap column (20 x 0.30 mm, 5 µm particle size; Phenomenex, Torrance, CA) over a period of 2 minutes at a flow rate of 10 µL/min using 100% solvent A. Peptides were eluted onto a Kinetex XB-C18 analytical column (150 x 0.30 mm, 2.6 µm particle size; Phenomenex) at a flow rate of 5 µL/min. Four µL of proteolytically digested peptides were loaded at 5% B and separated using a 45 minute linear gradient from 5 to 32% B, followed by an increase to 80% B for 1 minute, a hold at 80% B for 2 minutes, a decrease to 5% B for 1 minute, and a hold at 5% B for 6 minutes. The total HPLC acquisition time was 55 minutes. The following source parameters were used for all acquisitions: ion source gas 1 at 10 psi, ion source gas 2 at 25 psi, curtain gas at 30 psi, CAD gas at 7 psi, source temperature at 200°C, column temperature at 30°C, polarity set to positive, and spray voltage at 5000 V.

### Quantification by Data-Independent Acquisitions (SWATH DIA-MS)

For the data-independent acquisitions the survey MS1 spectra (m/z 395-1005) were acquired with an accumulation time of 100 ms, a declustering potential of 80 V and 0 V DP spread, and a collision energy of 10 V with 0 V CE spread (the time bins to sum were set to 8, and all channels were enabled). The MS2 spectra were collected within the same mass range as the MS1 spectra using 80 variable windows (**S1 Table**), with an accumulation time of 25 ms for each window, dynamic collision energy enabled, charge state 2 selected, and Zeno pulsing enabled. The total cycle time was 2.5 seconds.

### Data Processing and Quantification of SWATH DIA-MS (Spectronaut)

All data files were analyzed using Spectronaut (version 17.3.230224.55965; Biognosys) applying spectral library searches. The data base searches were conducted against the *Homo sapiens* pan human spectral library (31) with 20,526 entries, (32). Dynamic data extraction parameters and precision iRT calibration with local non-linear regression were used (33). Trypsin/P was specified as the digestion enzyme, allowing for specific cleavages and up to two missed cleavages. Methionine oxidation and protein N-terminus acetylation were set as dynamic modifications, while carbamidomethylation of cysteine was set as a static modification. Protein identification at the group level required at least 2 unique peptide identifications and was performed using a 1% q-value cutoff for both the precursor ion and protein level (34, 35). The iRT profiling was enabled. Protein quantification was based on the peak areas of extracted ion chromatograms (XICs) of 3 – 6 MS2 fragment ions with automatic local normalization and 1% q-value data filtering applied. The data was normalized using cross-acquisition normalizations.

### Further Data Processing and Visualizations

The ggplot2 package in R (version 4.0.5; RStudio, version 1.4.1106) were used for data visualization (36). Biological pathway analysis was performed using the Database for Annotation, Visualization, and Integrated Discovery (DAVID, accessed 06/25/2023) for over-representation analysis (ORA) of quantifiable proteins to determine which KEGG pathway gene ontology biological processes were enriched in these samples (37). Additional gene ontology analysis was performed using the Ingenuity Pathway Analysis (IPA, accessed 08/20/2023) tool of quantifiable proteins to determine which KEGG pathway gene ontology cellular and molecular functions were enriched in these samples (38).

### Data Availability

The raw data and complete mass spectrometry data sets have been uploaded to the Mass Spectrometry Interactive Virtual Environment (MassIVE) repository, which is maintained by the Center for Computational Mass Spectrometry at the University of California San Diego. These data can be accessed and downloaded using the following FTP link: ftp://MSV000098540@massive-ftp.ucsd.edu or via the MassIVE website: https://massive.ucsd.edu/ProteoSAFe/dataset.jsp?task=5a56838a5d804d48be8b2d0242319e70 (MassIVE ID: MSV000098540; ProteomeXchange ID: PXD066226).

## Results and Discussion

### Validation of iPSC-derived Neuron & Microglia Differentiation

iPSCs were differentiated towards neuronal and microglial lineages through a differentiation protocol that mimics in vivo developmental cues with specific growth factors and signaling molecules (**Figure 1A**). For neurons, iPSCs were differentiated into excitatory cortical neurons by forced expression of the transcription factor Neurogenin-2 (NGN2). As shown in **Figure 1B**, immunocytochemistry confirmed the successful differentiation into neurons by identifying key markers MAP2 (39), VGLUT1 for neurons (40). For microglia, a chemical induction protocol was used to differentiate iPSCs into microglia. The microglia were positive for Iba1, a known marker for microglia (41). These two methods generate physiologically relevant models for neuroscientific research and regenerative medicine.

**Figure 1.**
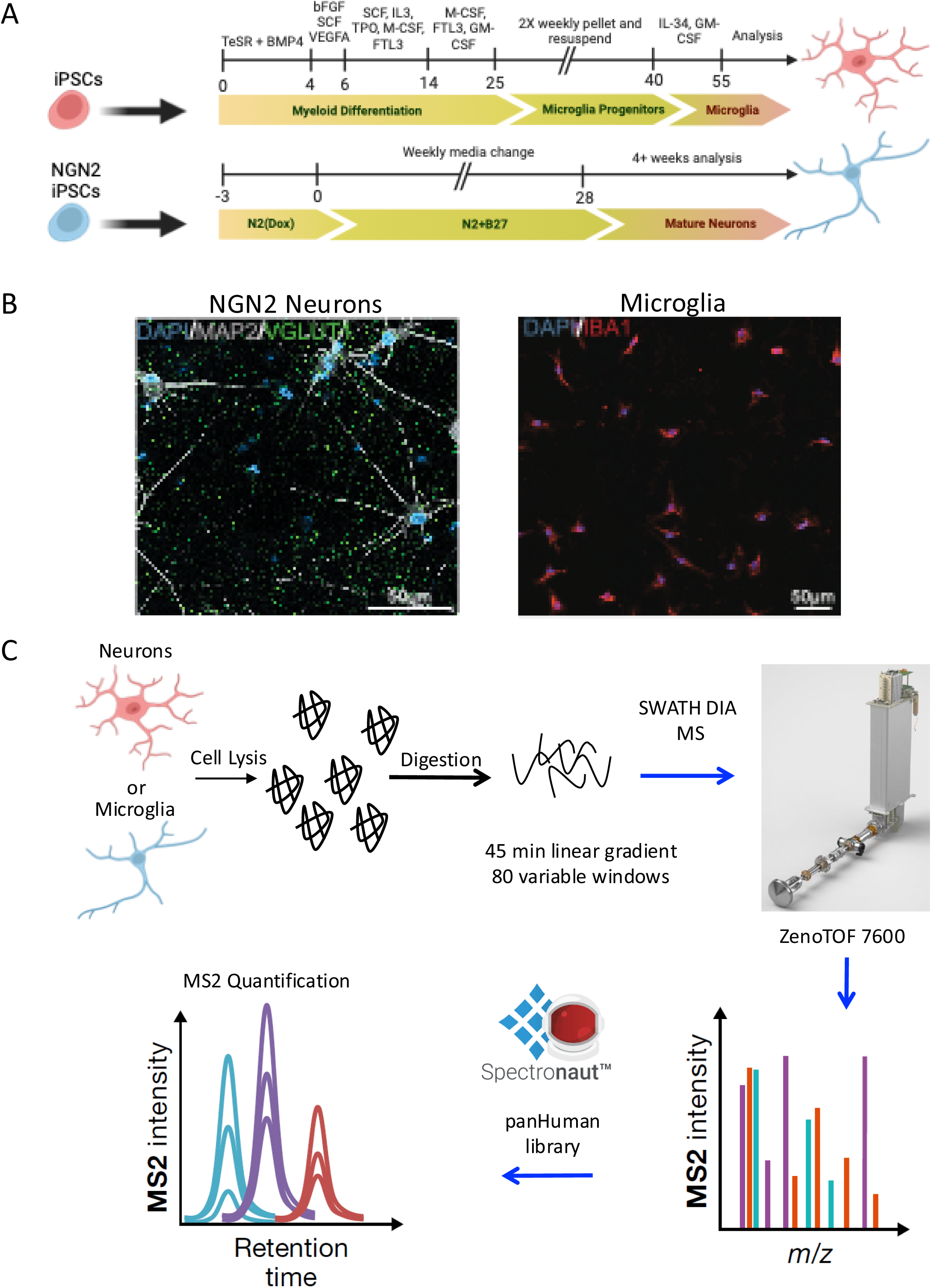
iPSC differentiation of microglia or neurons is confirmed through immunohistochemistry and proteomic analysis. A) Timeline for microglia (top) and neuron (bottom) differentiation from iPSCs. B) Staining of iPSC-Neurons and iPSC-Microglia. C) Overview of the proteomic workflow, including cell lysis, protein digestion, and analysis with data independent acquisition microflow chromatography and variable width windows (vw) using the ZenoTOF 7600 platform with the M-Class HPLC. DIA data was searched using the panHuman library in Spectronaut v17 for protein identification and quantification. R scripts were used for bioinformatic analysis.

### Proteomics Analysis of iPSC-derived Neurons and Microglia – Protein Signatures

To fully characterize each brain cell type we carried out a proteomics analysis. We acquired DIA-MS data for proteolytical digestions of iPSC-derived microglia (n = 6) and iPSC-derived neurons (n = 6), respectively, each in technical triplicate, utilizing a 45-minute microflow chromatographic gradient (**Figure 1C**). We carried out separate searches for the neuron and microglia data using Spectronaut v17. This analysis resulted in the identification of 4,178 unique protein groups quantified with 2 or more unique peptides in microglia samples (**S2 Table**) and 4,484 unique protein groups quantified with 2 or more unique peptides in neuronal samples (**S3 Table**, **Figure 2A,B**). The proteomic profiles overlapped approximately 80% with 3,347 quantified protein groups commonly found in both microglia and neuronal lysates (**Figure 2C**). However, we also quantified 817 protein groups identified uniquely in iPSC-derived microglia and 1,130 protein groups uniquely identified in iPSC-derived neurons, underscoring the distinct cellular functions of neurons and microglia.

**Figure 2.**
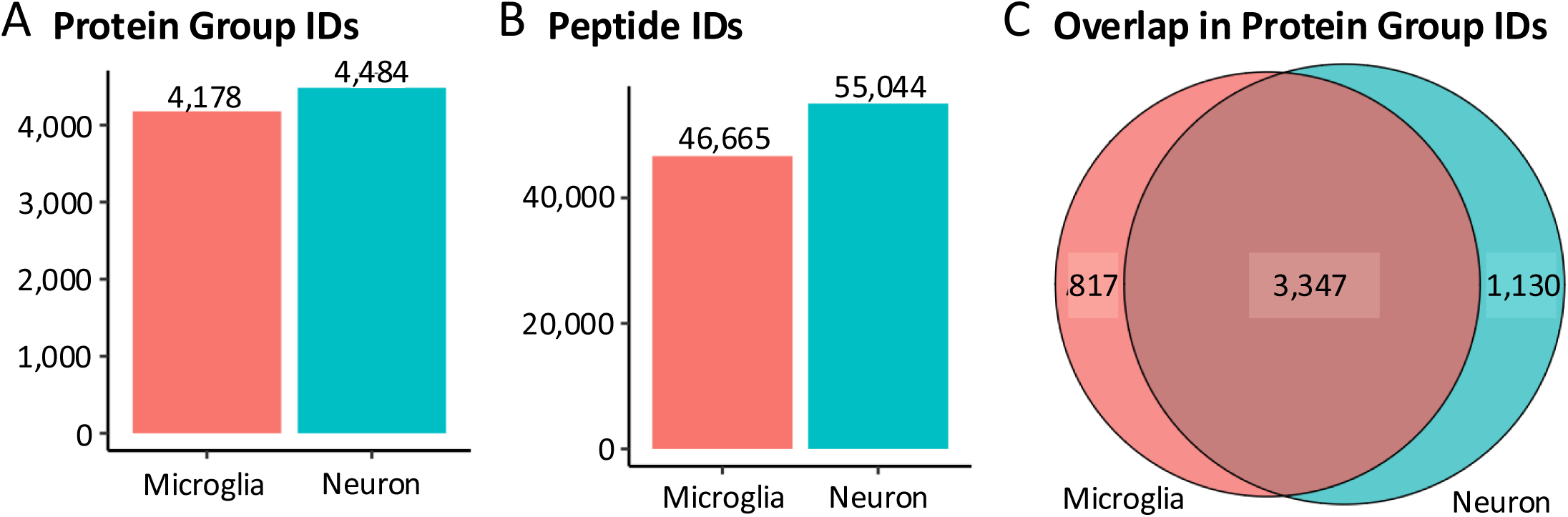
Reproducible identification of microglia and neuronal cell lysates on a 45-min linear microflow chromatography gradient. A) Protein group and B) peptide identifications (IDs) for each cell line, and F) the overlap in Protein Group IDs is shown for each cell line.

Our mass spectrometric analysis revealed distinct sets of protein markers specific to neurons and microglia (39, 42), as well as subsets of proteins that were common to both cell types (43) (see **Figure 3A**). For neurons, markers, such as Amyloid Precursor Protein (APP), Calbindin 1 (CALB1), Calretenin (CALB2), Disks Large MAGUK Scaffold Protein (DLG) 1, DLG3, DLG5, Growth Associated Protein 43 (GAP43), Neurofilament Light Chain (NEFL), Microtubule-Associated Protein 2 (MAP2) and MAP Tau (MAPT) are indicative of synaptic function and neuronal structure (44). These markers are involved in processes, such as synaptic plasticity, neuroprotection, and microtubule stabilization, and are key to understanding neuronal behavior and pathology, especially in neurodegenerative diseases like Alzheimer’s disease. In contrast, microglia-specific markers like Allograft Inflammatory Factor 1 (AIF1), Monocyte Differentiation Antigen CD14, Scavenger Receptor Cysteine-rich Type 1 Protein M130 (CD163), Tumor Necrosis Factor Receptor Superfamily Member 5 (CD40), Matrix Metalloproteinase-9 (MMP9), and Integrin Alpha-M (ITGAM) reflect the roles in immune response and phagocytosis (45). These proteins are integral to the microglia function as the resident immune cells in the brain, responding to injury and disease by mediating inflammatory responses and facilitating tissue repair. Shared markers, including Enolase 2 (ENO2), Tubulin Beta Class III (TUBB3), and Vimentin (VIM) suggest essential roles in basic cellular processes and cytoskeletal organization, underscoring the interconnectedness of cellular functions in the brain’s microenvironment (46) (47). The differentiation process clearly shows the cellular specificity and functional distinctiveness of neurons and microglia derived from iPSCs.

**Figure 3.**
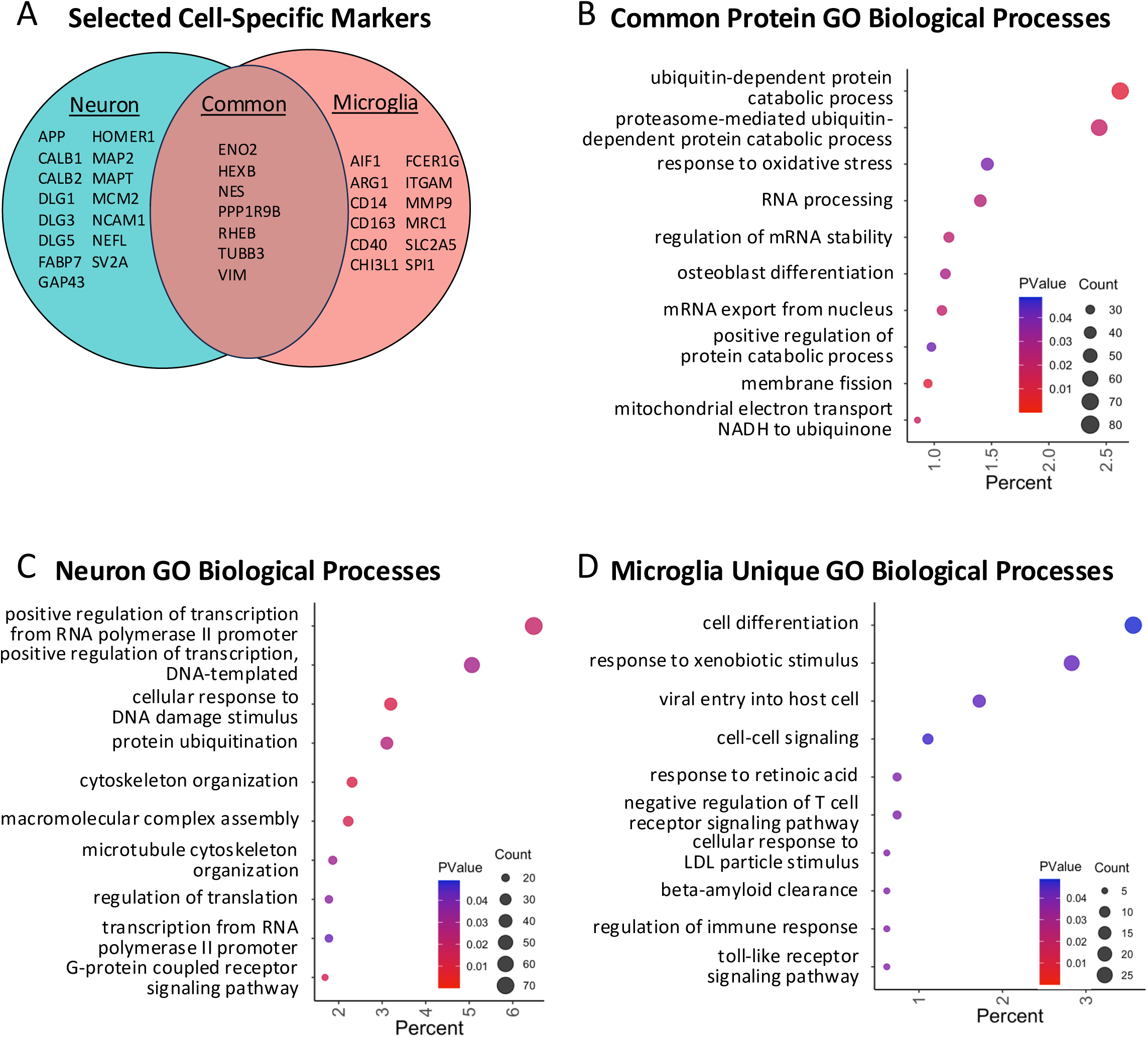
Cell-specific markers and biological processes are identified in microglia and neuronal cell lysates. A) The cell-specific neuron and microglia protein markers identified in each dataset or in common are shown in a Venn diagram. Gene Ontology (GO) biological processes are shown for proteins B) common to both data sets, C) unique to neuron lysates, or D) unique to microglia lysates. GO terms, identified with the DAVID pathway analysis tool (37), are filtered with a p-value ≤ 0.05 cut-off.

Gene Ontology (GO) biological processes were identified for unique and shared proteins in the microglia and neuron data using DAVID: Database for Annotation, Visualization, and Integrated Discovery (37). The GO biological processes common to both cell types (**Figure 3B**) suggested that several fundamental cellular functions were conserved across neurons and microglia. These processes included ubiquitin-dependent protein catabolic process, response to oxidative stress, and regulation of mRNA stability, among others. These shared pathways highlighted the universal cellular responses to environmental stimuli and internal regulatory mechanisms that are critical for maintaining cellular homeostasis. Unique to the investigated neuron populations were specialized functions, such as synaptic signaling and neurogenesis (**Figure 3C**). The prominence of processes related to synaptic organization and neuronal communication underscored the neuronal pivotal roles in information processing and transmission within the nervous system. These findings aligned with the known physiological roles of neurons and further elucidated their specific contributions to neural network dynamics and brain function. The GO biological processes for microglia (**Figure 3D**) emphasized the role of microglia in immune response, phagocytosis, and inflammatory signaling. The identification of these processes emphasized the role of microglia as the resident immune cells of the brain, involved in pathogen defense, debris clearance, and regulation of inflammatory responses. Our results are consistent with the critical role of microglia in maintaining brain health and their involvement in neuroinflammatory and neurodegenerative diseases.

### Delineating Neuronal and Microglial Roles in Neuroinflammatory Pathways

To further explore the cellular and molecular functions associated with the human iPSC-derived neuron and microglia proteomes, we performed an Ingenuity Pathway Analysis (38). The data presented in **Figure 4A** shows the distinct molecular signatures and functional activities of neurons and microglia within the central nervous system. Notably, the analysis revealed significant activation differences between neurons and microglia. Neuron-specific pathways such as Neuregulin Signaling and Reelin Signaling in Neurons highlight critical processes in synaptic function and neuronal plasticity. The Synaptic Long-Term Potentiation pathway, activated predominantly in neurons, underscores the fundamental mechanisms underpinning learning and memory. Conversely, the prominence of Neuroinflammation Signaling in microglia delineates their pivotal role in neuroinflammatory processes, which can contribute to neurodegenerative conditions. The differential expression of IL-10 and IL-12 signaling pathways, primarily in microglia, emphasizes the role of these cells in modulating inflammatory responses, with IL-10 signaling associated with anti-inflammatory effects and IL-12 with pro-inflammatory responses. The activation of the Acute Phase Response Signaling and Macrophage Classical Activation pathways in microglia further supports their role in the innate immune response within the brain. This comparative analysis elucidates the complex interplay between neurons and microglia in maintaining neural homeostasis and responding to pathological states. Understanding these distinct cellular and molecular functions is crucial for developing targeted therapeutic strategies aimed at modulating neuroinflammatory and neurodegenerative processes.

**Figure 4.**
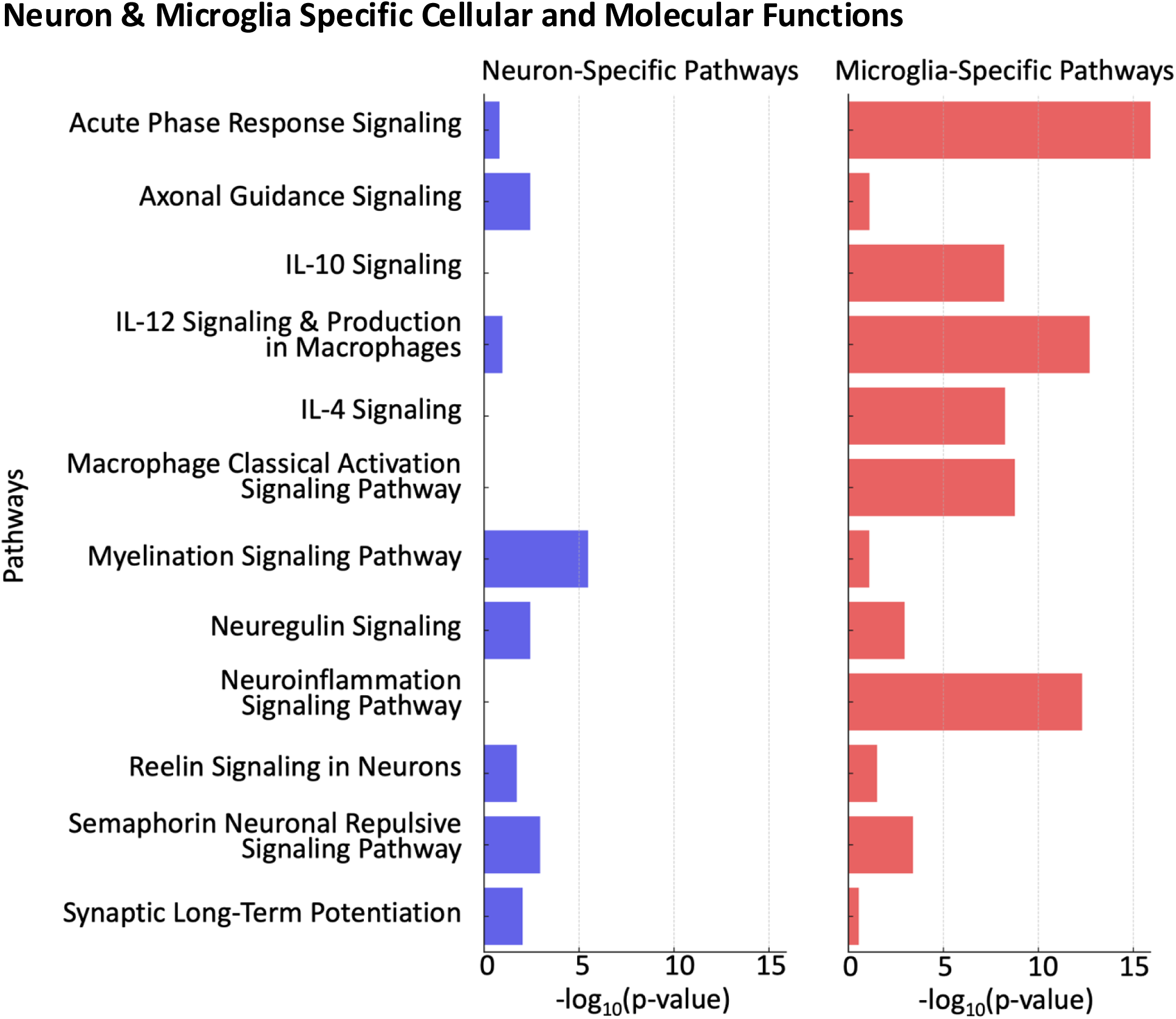
Comparative analysis of neuron and microglia specific cellular and molecular functions. The histogram illustrates the differential activation of cellular and molecular pathways in neurons and microglia identified with the Ingenuity Pathway Analysis tool (38), as evidenced by the -log10(p-value) of various signaling pathways, including Acute Phase Response, Axonal Guidance, IL-10, IL-12 in Macrophages, IL-4, Macrophage Classical Activation, Myelination, Neuregulin, Neuroinflammation, Reelin in Neurons, Semaphorin Neuronal Repulsive, and Synaptic Long-Term Potentiation. The pathways are divided into neuron-specific and microglia-specific categories, highlighting the distinct functional roles each cell type plays in the context of neuroinflammation and neurodegeneration.

### Step by Step Protocol of Data-Independent Acquisitions on the ZenoTOF 7600

This study presents a comprehensive tutorial on optimizing data-independent acquisition (DIA) for the quantitative proteomic analysis of iPSC-derived neurons and microglia, utilizing the advanced capabilities of the SCIEX ZenoTOF 7600 mass spectrometer in conjunction with a Waters M-Class LC system (**Supplementary Protocol Methods**). The protocol demonstrates the development of an optimized 45-minute gradient DIA method tailored for the ZenoTOF 7600 system coupled to the M-Class HPLC. Key acquisition parameters included an MS1 accumulation time of 100 ms, MS2 accumulation time of 25 ms, charge state 2 selection, and enabling Zeno pulsing. For data processing, the pan-Human library in Spectronaut v17 was used with library-free searching in directDIA mode. This tutorial provides a framework to optimize DIA-MS methods on the ZenoTOF 7600 system to sensitively and reproducibly quantify proteomes of cell models, organoids, animal tissue samples, patient (disease) samples, and other scarce sample types that are critical for many biological and clinical studies.

## Limitations of this Study

While this study presents a robust protocol for iPSC differentiation and an optimized DIA SWATH MS method for deep proteomic profiling, the use of monocultures, though essential for clear proteomic characterization of individual cell types, does not capture the dynamic interactions that occur between these cells in vivo. Recent studies have demonstrated that microglia exhibit a more homeostatic state co-cultured with neurons, including lower pro-inflammatory cytokine secretion and changes in ramification patterns (48, 49). Similarly, neurons show altered morphology, and modified network activity in the presence of microglia due to their role in synapse regulation and debris clearance (ref). Therefore, while the current approach provides baseline data, future studies incorporating co-culture systems and organoid models would be needed to understand how cell-cell interactions alter proteomic signatures and better recapitulate in vivo neuroglial dynamics. Additionally, comparative analyses with in vivo samples would further validate the translational relevance of these findings.

This study focused on developing a robust mass spectrometric assay and protocol to easily monitor and quantify the proteomes of iPSC-derived neurons and microglia. The ultimate goal will be to apply this workflow to study disease-relevant cohorts and therapeutic interventions. Future studies will include iPSC lines derived from patients with neurodegenerative diseases to assess the ability of the optimized SWATH DIA method to detect and quantify disease-associated proteomic changes. Additionally, the incorporation of other omics data, such as transcriptomics and metabolomics, could provide a more comprehensive view of the cellular mechanisms at play. Finally, the bioinformatic analyses performed here, do not yet investigate the detailed mechanisms or functional consequences of the identified proteomic differences between iPSC-derived neurons and microglia. Follow-up experiments, such as targeted protein quantification, pathway perturbation, and functional assays, will be necessary to validate and extend the findings of this study.

## Conclusion

We presented a comprehensive tutorial on optimizing data-independent acquisition (DIA) for the quantitative proteomic analysis of iPSC-derived neurons and microglia, utilizing the advanced capabilities of the SCIEX ZenoTOF 7600 mass spectrometer coupled to a Waters M-Class LC system. Our study not only showcases the methodological advancements in mass spectrometry-based proteomics but also highlights its application in understanding the proteomic landscapes of cell models relevant to neurodegenerative disease research. By leveraging the unique features of the ZenoTOF 7600 system we achieved a significant enhancement in the duty cycle and sensitivity, facilitating the identification and quantification of thousands of proteins.

Our findings provide valuable insights into the distinct and overlapping proteomic signatures of neurons and microglia, shedding light on their specific roles in the central nervous system. The identification of cell-type-specific markers and the elucidation of their functional pathways underscore the utility of iPSC-derived cell models in mimicking the in vivo environment and advancing our understanding of neurobiology. The comparative proteomics approach employed in this study highlights the power of DIA-MS in revealing the complexity of the proteome and its dynamic changes between cell types, offering a pathway forward for deeper understanding of the molecular basis of neurodegenerative diseases. Furthermore, the validation of the iPSC differentiation protocols, as evidenced by the successful identification of key neuronal and microglial markers, establishes a reliable framework for generating physiologically relevant models of the human brain. These models are indispensable for exploring the underlying mechanisms of neurodegeneration and for screening potential therapeutic interventions.

The combination of robust iPSC differentiation and sensitive proteomic characterization described here provides a valuable platform for future studies investigating the cellular mechanisms underlying neurological disorders. Quantifying dysregulated proteins and pathways in patient-derived neurons and microglia could provide insights into disease etiology and progression. Moreover, this DIA approach enables deep proteome profiling of precious iPSC-derived cell models, enhancing their utility for investigating disease biology and therapeutic development.

## Supporting information

Supplementary Protocol Method

## Acknowledgements

The authors acknowledge the very generous support from SCIEX for the ZenoTOF 7600 system and a Waters M-class HPLC system at the Buck Institute.

## Authorship Contribution

BS and LME: Conceptualization, Acquisition of Funding, Supervision, editing of manuscript; JBB: data acquisition; TM: sample generation, Editing of manuscript; CAS: data processing, editing of manuscript; JB: data processing, editing of manuscript.

## Supplemental Material

**S1 Table. ZenoTOF 7600 data-independent acquisition isolation window scheme.**

The 80 variable width window isolation scheme used for data-independent acquisition on a 45-minute microflow gradient (5 µL/min) is shown.

**S2 Table. Proteins identified searching the DIA-MS microglia data using a pan-Human spectral library.**

Data for the 4,178 protein groups identified with 2 or more unique peptides from microglia cell lysates using Spectronaut v17 using the Pan-Human spectral library published by Rosenberger et al. (32)

**S3 Table. Proteins identified searching the DIA-MS microglia data using a pan-Human spectral library.**

Data for the 4,484 protein groups identified with 2 or more unique peptides from neuron cell lysates using Spectronaut v17 using the Pan-Human spectral library published by Rosenberger et al. (32)

**Supplementary Tutorial Method. Generation of a data-independent acquisition method for a SCIEX quadrupole time of flight ZenoTOF 7600 platform, and instructions for library-free data processing with the panHuman library or directDIA in Spectronaut.**

This tutorial presents an optimized DIA method tailored for a SCIEX ZenoTOF 7600 mass spectrometer coupled to a Waters M-Class liquid chromatography system. The optimized method was used to acquire DIA data from tryptic digests of iPSC-derived neurons and microglia. The peptide search space was established using the library-free directDIA algorithm in Spectronaut v17. The DIA parameters presented here provided the optimal balance of quantification and depth of detection for this specific instrument configuration and sample type. By following this tutorial, tailored DIA methods can be developed that fully leverage the capabilities of the mass spectrometer system.

