## Supplementary Protocol Method for "Investigation of Neurons and Microglia from *APOE3* Induced Pluripotent Stem Cells using Data-Independent Acquisitions on a ZenoTOF 7600"

**Supplemental Protocol**

**Variable Window Data-independent Acquisition Method Setup for a ZenoTOF 7600 Mass Spectrometer, and Instructions for Data Processing with directDIA (Spectronaut)**

Note: use Hyperlinks to go to the individual chapters by clicking on the page numbers above.

### **Chapter 1. Introduction to the SCIEX ZenoTOF 7600 and Waters M-Class Platform**

This tutorial is for the setup, collection, processing, and report generation of bottom-up proteomics data-independent acquisition data collected on a SCIEX ZenoTOF 7600 mass spectrometer coupled to a Waters M-Class liquid chromatography system with a 1- 10 µL electrode in an OptiFlow® Turbo V Ion Source and processed using Spectronaut v17. The Waters M-Class liquid chromatography system is operated with a Trap-Elute configuration using a Phenomenex Luna® 5 µm C18 100 A Micro Trap (20 x 30 mm) and a Phenomenex Kinetex® 2.6 µm XBV-C18 100 A LC Column (150 x 0.3 mm). The trap flow rate is 1 µL/min and the analytical flow rate is 5 µL/min.

### **Chapter 2. Introduction to SCIEX OS**

1. Open **SCIEX OS**.

The **SCIEX OS** homepage allows access to multiple **Acquisition, Processing,** and **Management** Perspectives. These perspectives can also be accessed by clicking the top left icon. The right side of the homepage shows the status of the activated and operating Devices.

*Note: Here we use SCIEX OS version 3.1.0.16485.*

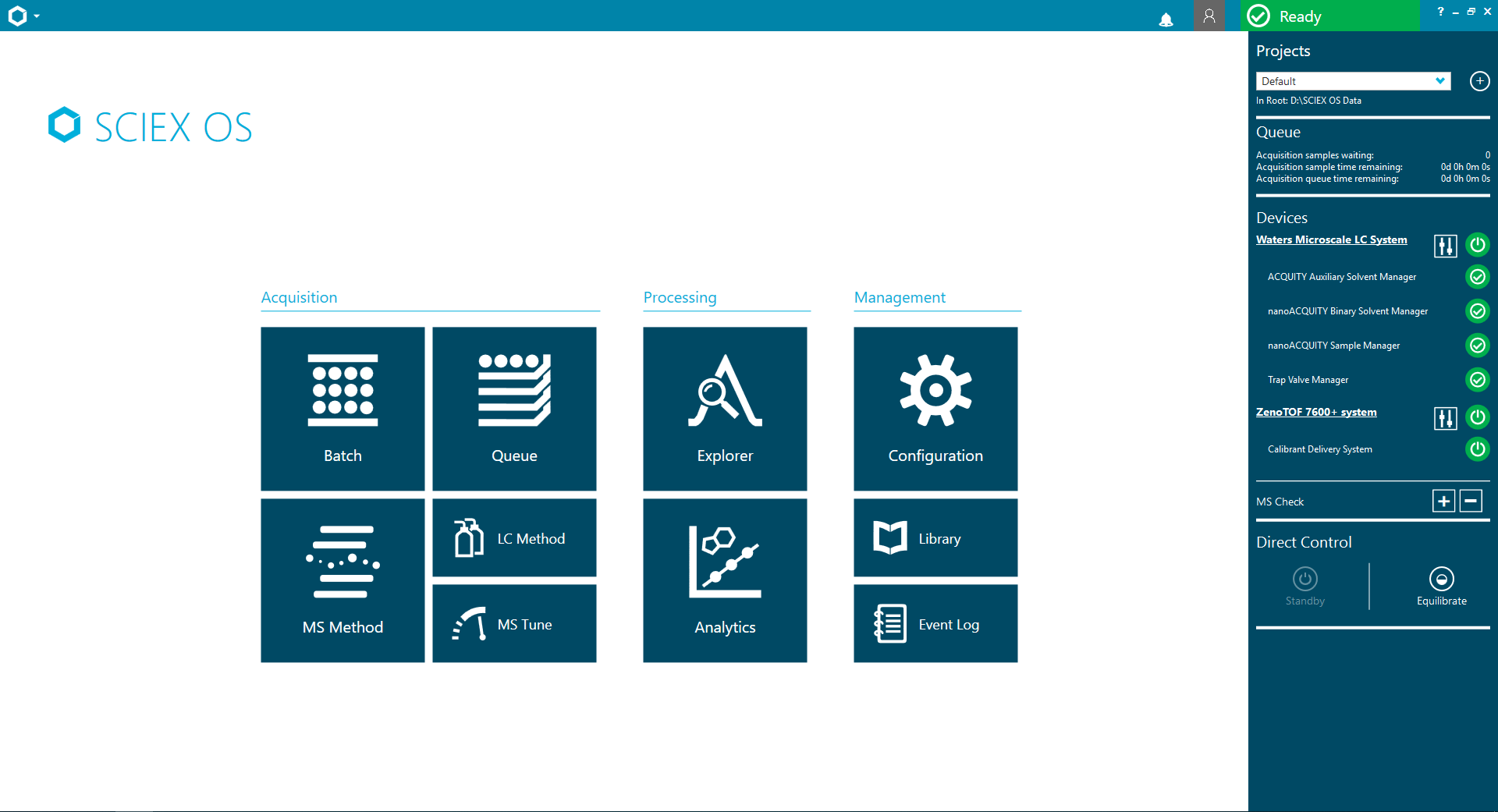

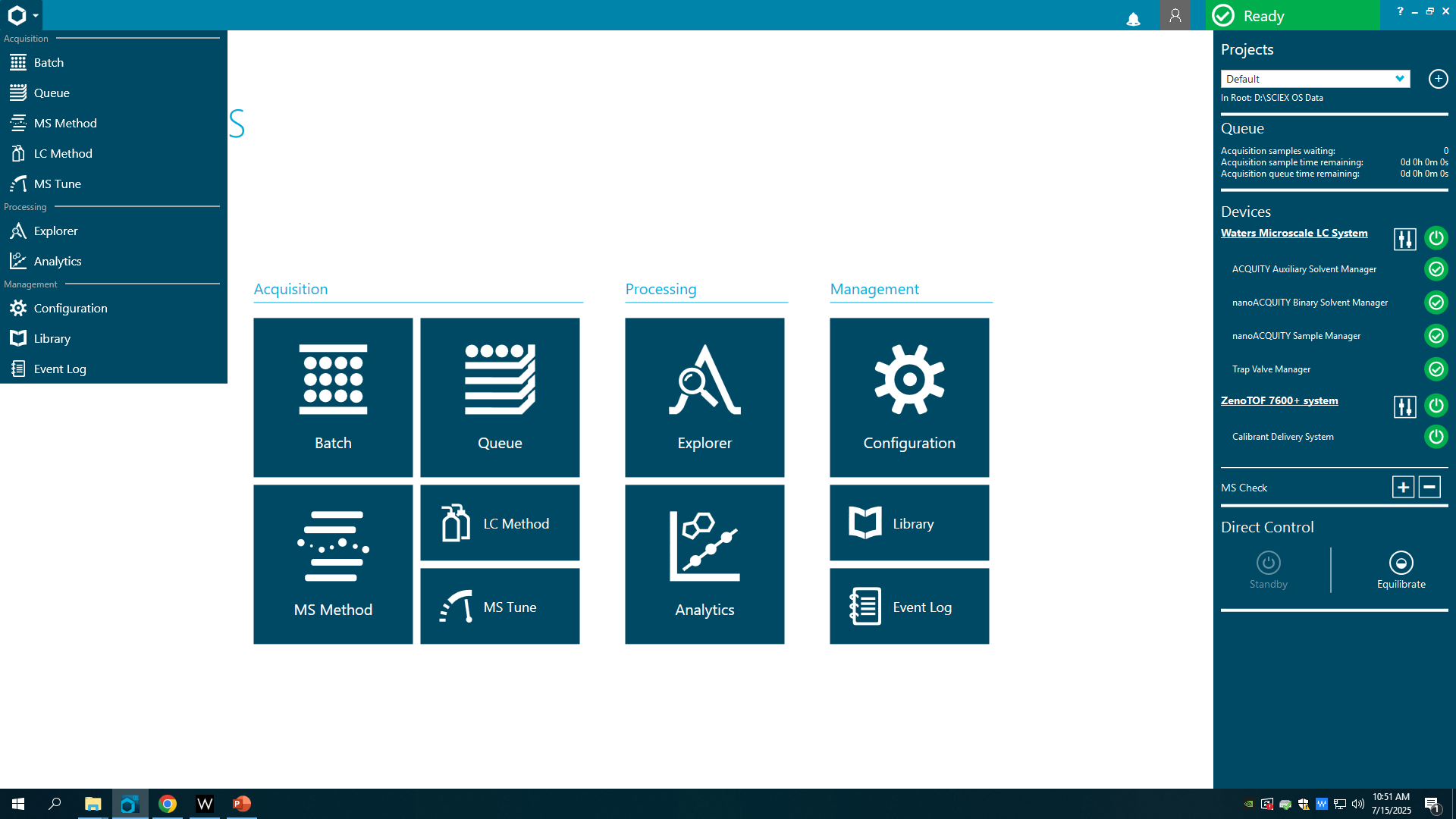

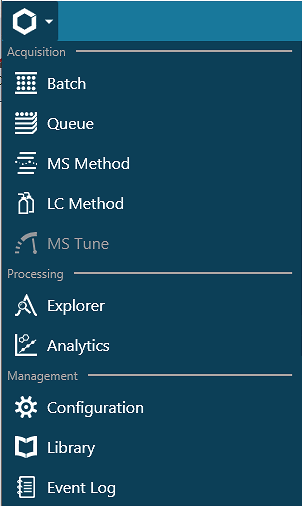

Perspectives:

- **Batch**
  - Set up sample acquisition and data processing
- **Queue**
  - Monitor sample acquisition
- **MS Method**
  - Set up mass spectrometry methods including data dependent acquisition (IDA), data independent acquisition (SWATH), and multiple reaction monitoring (MRM HR and Guided MRM HR).
- **LC Method**
  - Set up liquid chromatography methods
- **MS Tune**
  - Tune and assess the performance of the mass spectrometer
- **Explorer**
  - Visualize and assess quality of data
- **Analytics**
  - Analyze data with SCIEX OS
- **Configuration**
  - Activate devices and set SCIEX OS parameters
- **Library**
  - Compound libraries for data analysis using SCIEX OS
- **Event Log**
  - Log of errors and acquisition events through data acquisition and processing

2. Open the **Configuration** perspective and ensure that devices are activated. If they are not active, they can be activated by checking the right box and clicking activate.

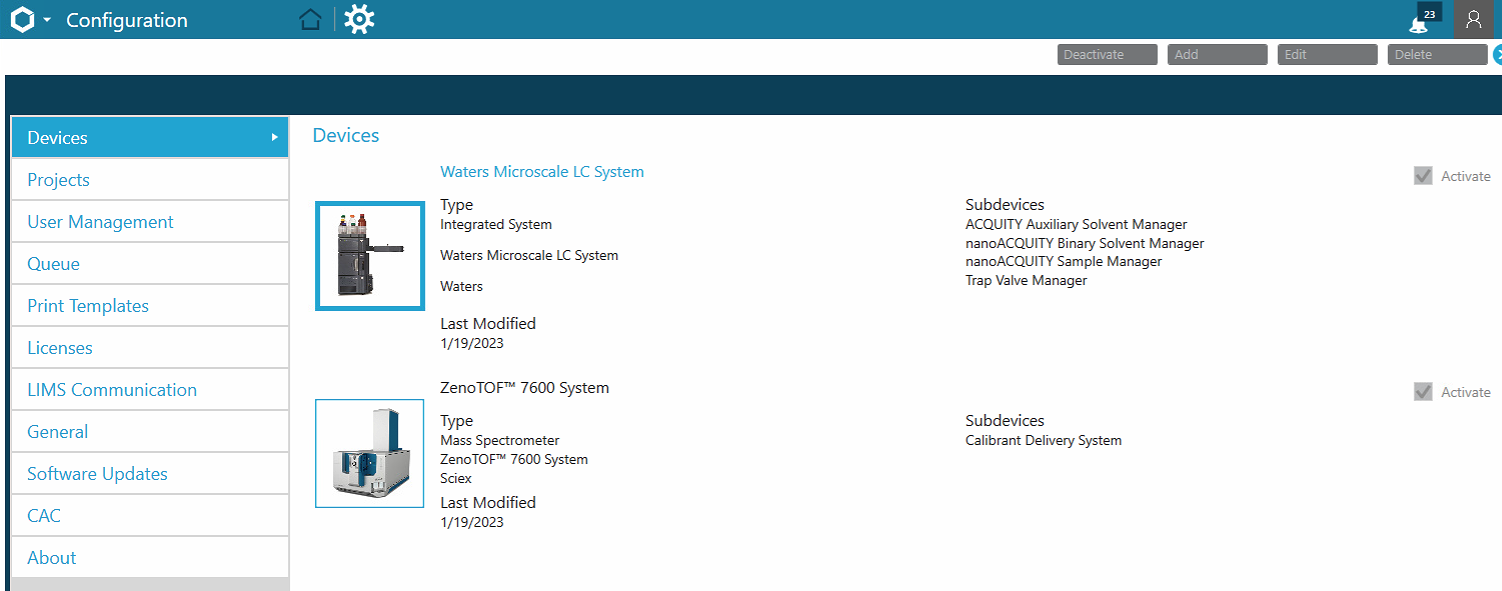

### **Chapter 3. Generate a Mass Spectrometry method file for variable window ZenoSWATH Data Independent Acquisition**

1. Navigate to the **MS Method** perspective.

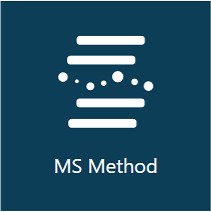

2. Click **New** and select **SWATH** to start a new method.

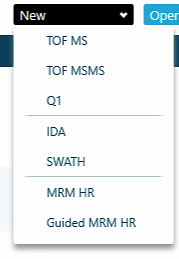

The **MS Method** Perspective:

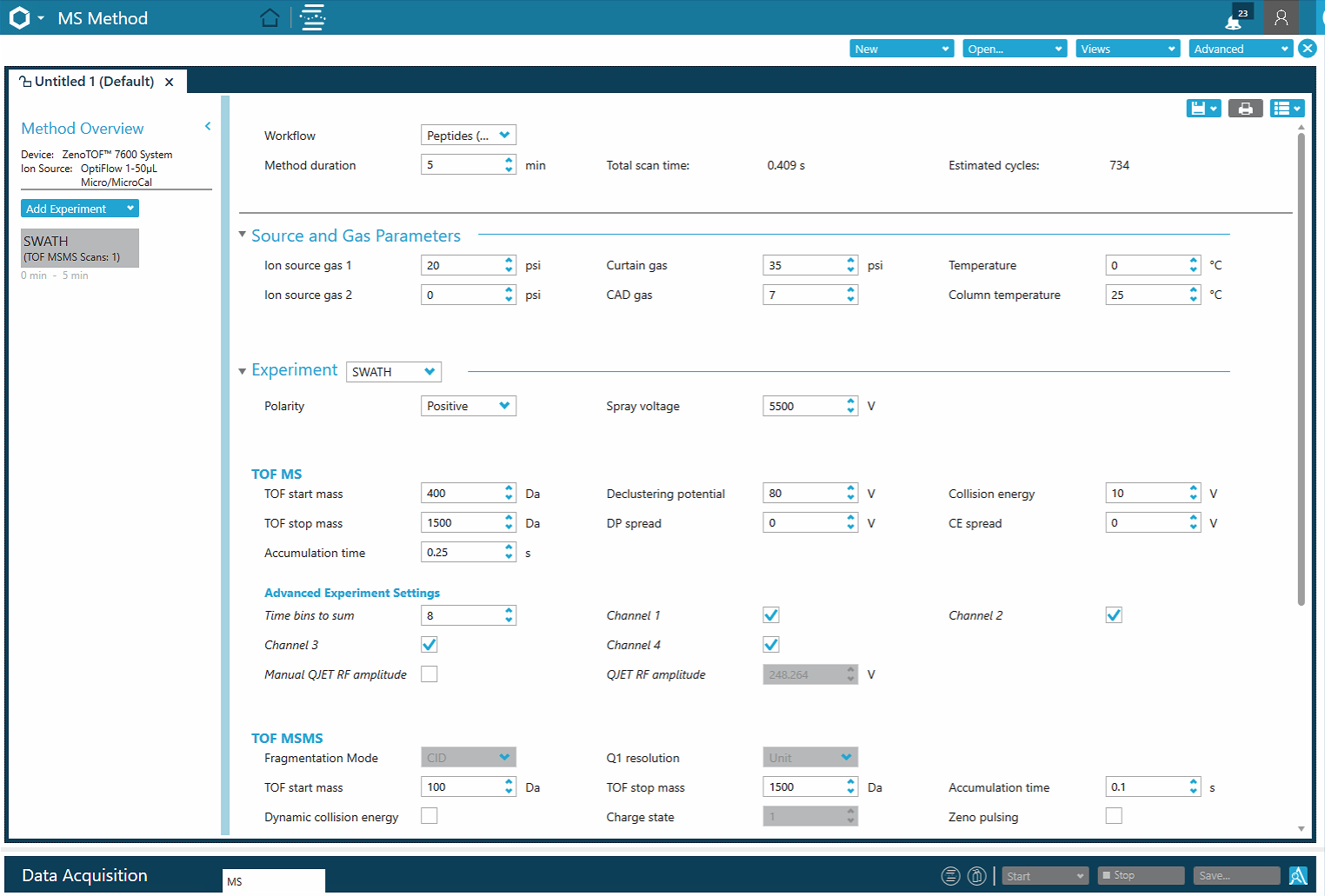

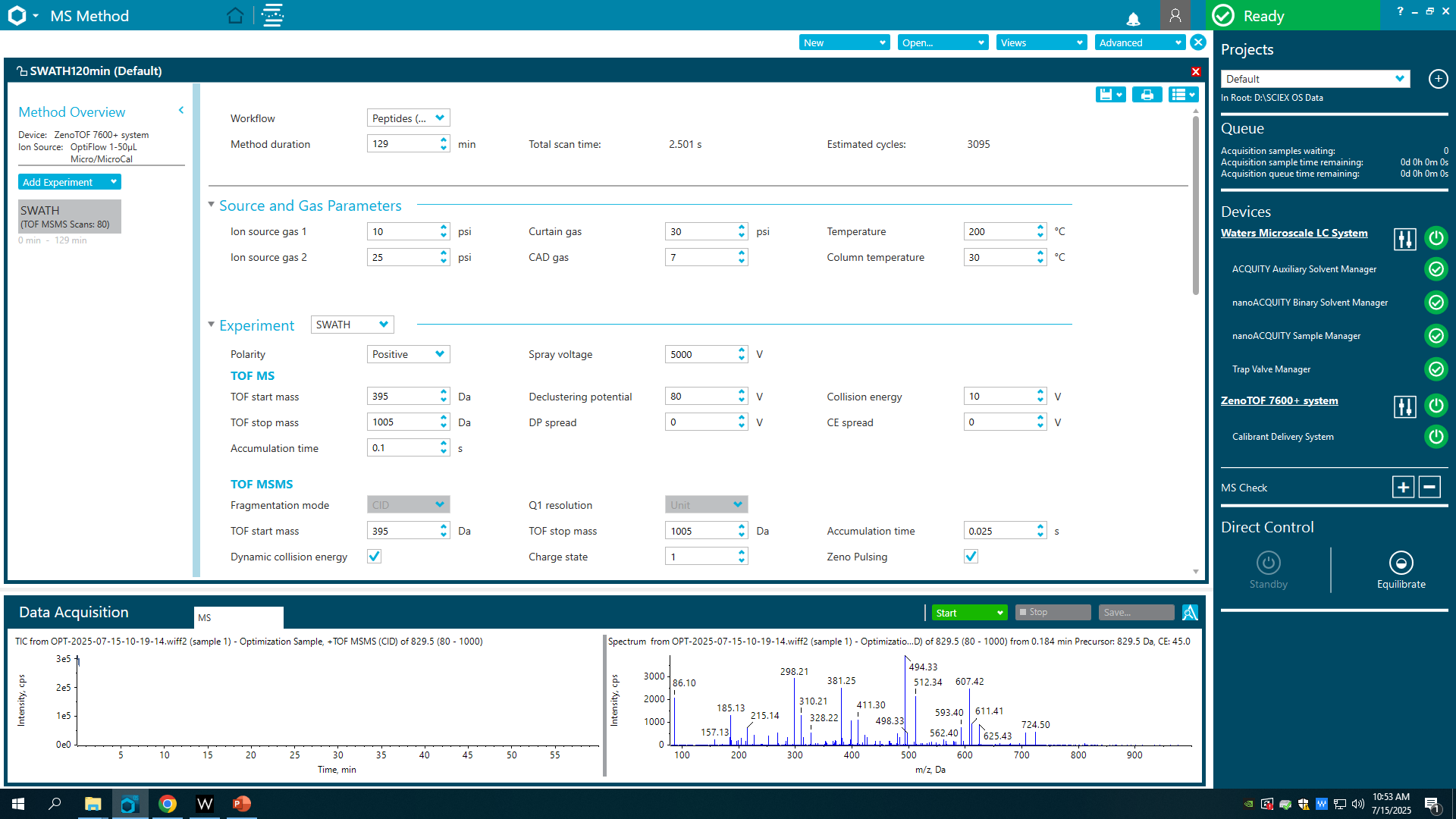

3. Click the **Workflow** dropdown and select **Peptides (or other ≥2± analytes with MW ≤ 5kDa)**.

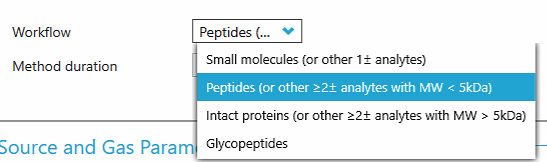

4. Set the **Method Duration** to the length of the chromatographic gradient.

*Note: In this study, the length of the chromatographic gradient is 129 min.*

5. On the **Source and Gas Parameters** menu, change the following parameters:

- set the **Ion source gas 1** to 10 psi
- set the **Ion source gas 2** to 25 psi
- set the **Curtain gas** to 30 psi
- set the **CAD gas** to 7
- set the **Temperature** to 200 °C
- set the **Column temperature** to 30 °C.

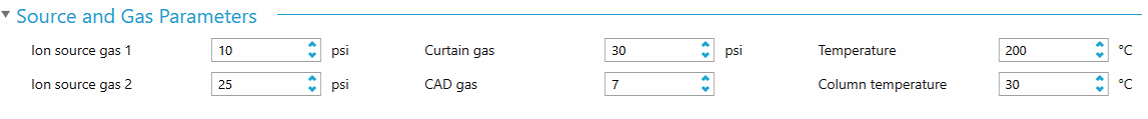

6. On the **Experiment** menu:

- set the **Experiment** to SWATH
- set the **Polarity** to Positive
- set the **Spray voltage** to 5000.

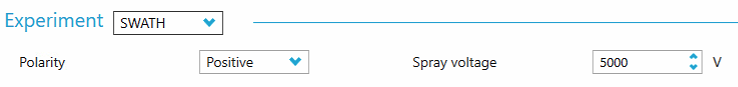

7. Set the **TOF MS settings:**

- set the **TOF start mass** to 395 Da
- set the **TOF stop mass** to 1005 Da
- set the **Accumulation time** to 0.1s
- set the **Declustering potential** to 80 V
- set the **DP spread** to 0 V
- set the **Collision energy** to 10 V
- set the **CE spread** to 0 V.

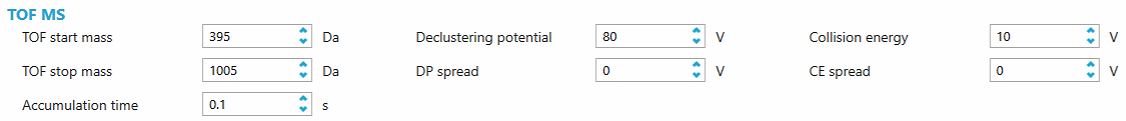

8. Under the **TOF MS settings,** set the **Advanced Experiment Settings:**

- set the **Time bins to sum** to 8
- ensure that **Channel 1, Channel 2, Channel 3,** and **Channel 4** are enabled
- ensure the **Manual QJET RF amplitude** is disabled.

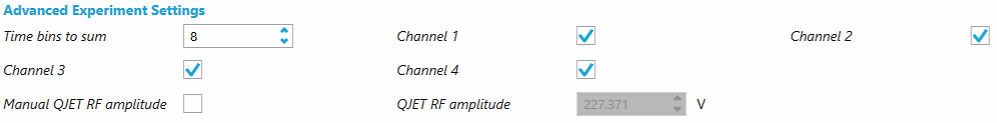

9. On the **TOF MSMS**, change the following parameters:

- ensure the **Fragmentation Mode** is set to CID
- set the **TOF start mass** to 395 Da
- enable the **Dynamic collision energy**
- ensure the **Q1 resolution** is set to Unit
- set the **TOF stop mass** to 1005 Da
- set the **Charge state** to 2
- set the **Accumulation time** to 0.025 s
- enable **Zeno Pulsing.**

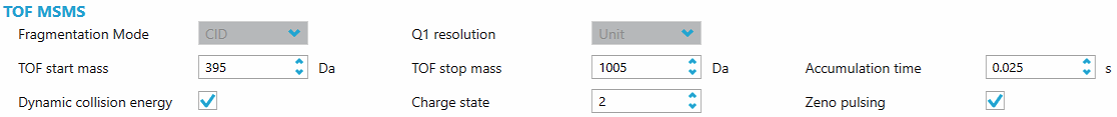

10. Under the **TOF MSMS settings,** set the **Advanced Experiment Settings:**

- set the **Time bins to sum** to 8
- ensure that **Channel 1, Channel 2, Channel 3,** and **Channel 4** are enabled
- ensure the **Manual QJET RF amplitude** is disabled.

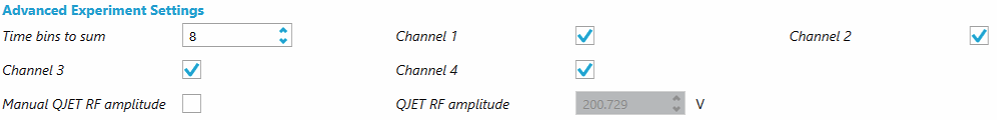

11. Import the isolation window strategy to the **TOF MSMS Mass Table**

- prepare the **DIA Variable Window text file** from the Supplementary Table S1 that defines the variable width isolation window strategy
- copy and paste the table from the Supplementary Table S1 to the **Mass table**
- disable and re-enable the **TOF MSMS Dynamic collision energy** setting to populate the collision energies into the **Mass Table**

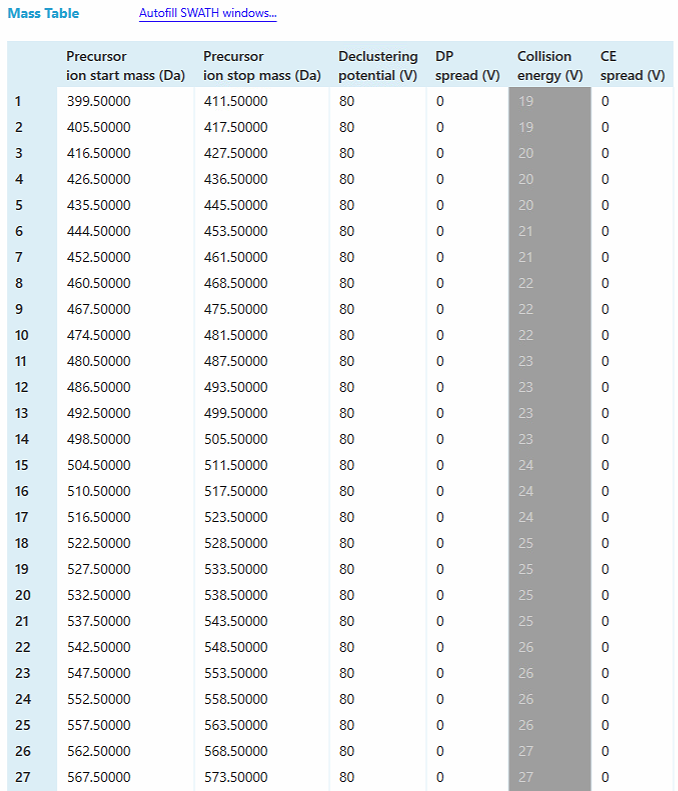

12. Click **Floppy Disk Icon** and **Save** the instrument method file.

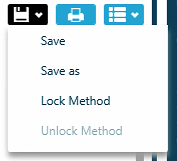

### **Chapter 4. Generate a Liquid Chromatography Method File in SCIEX OS**

1. Open the **LC Method** perspective.

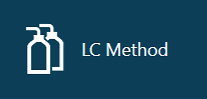

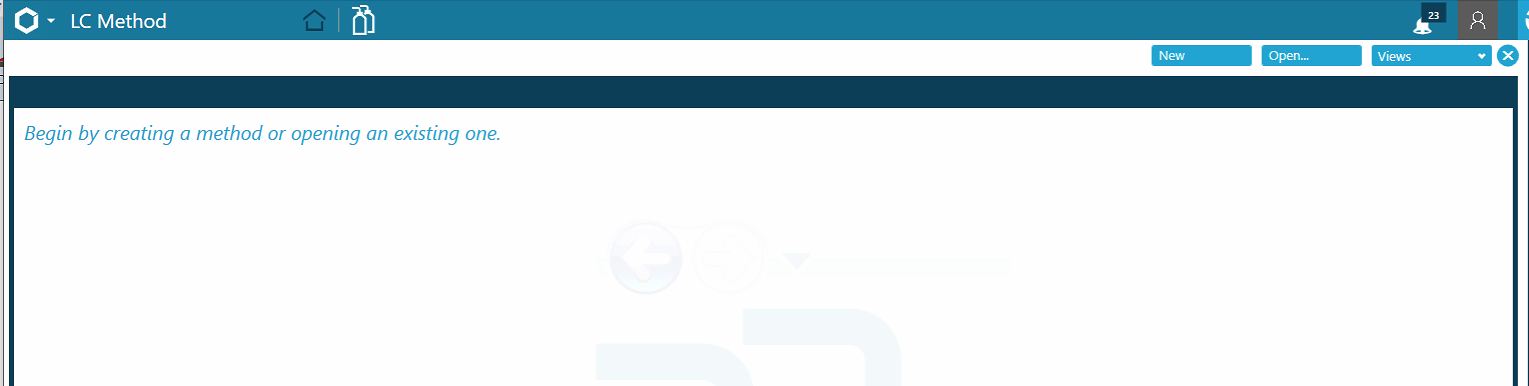

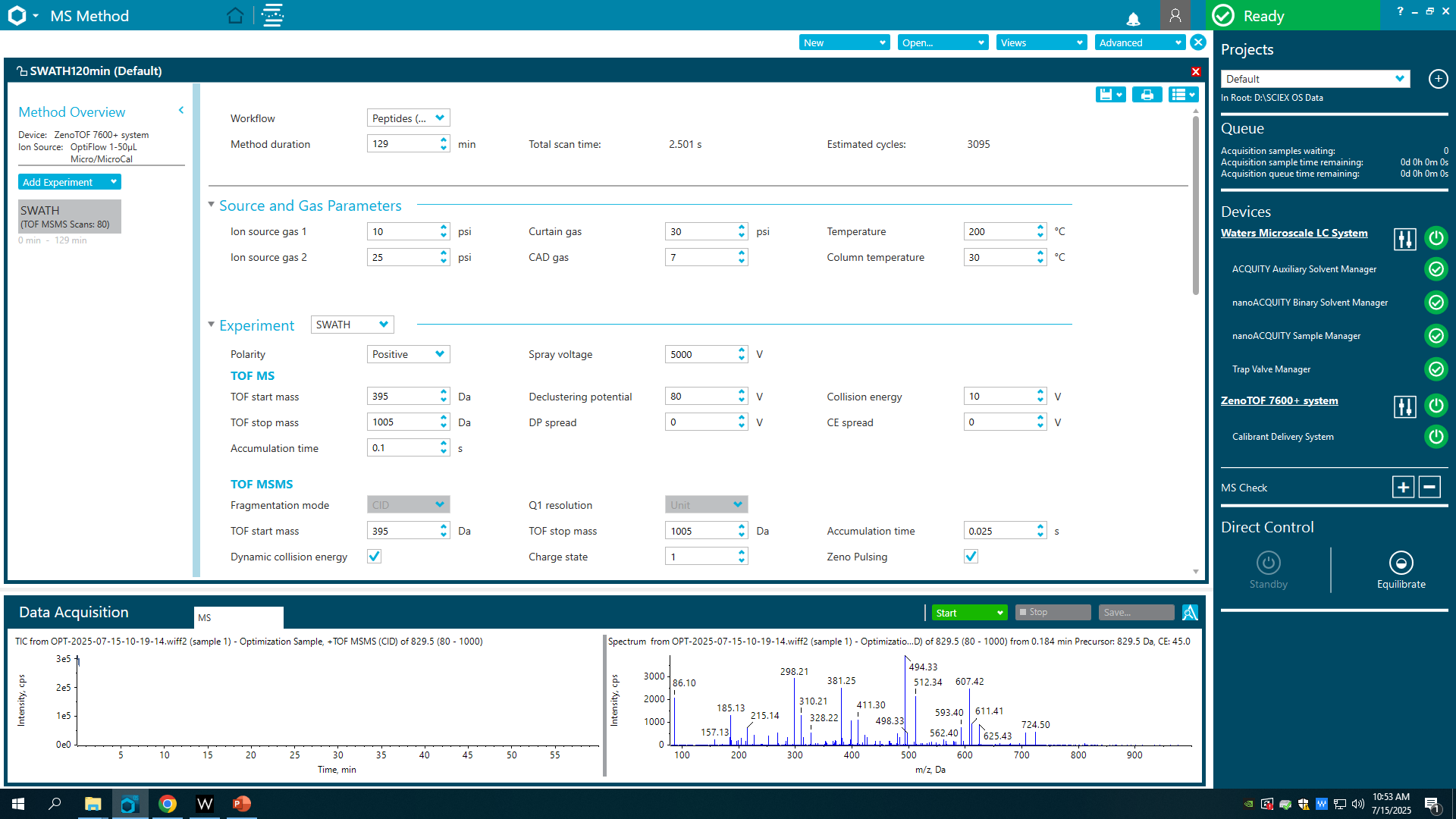

2. Click **New** to set up a new LC Method.

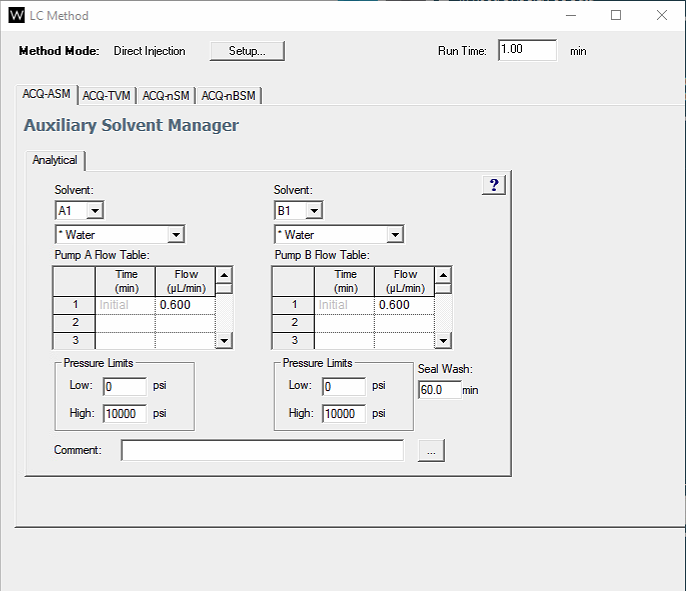

3. Click the **Setup** button to change the **Method Mode** to “Trapping”:

- Set the **Loading Time** to 3 min
- Ensure that “Disable Multi-Load” is selected
- Ensure that “Flow rate is ramped to zero, valve position changes, flow rate is ramped” is selected
- Click **OK**.

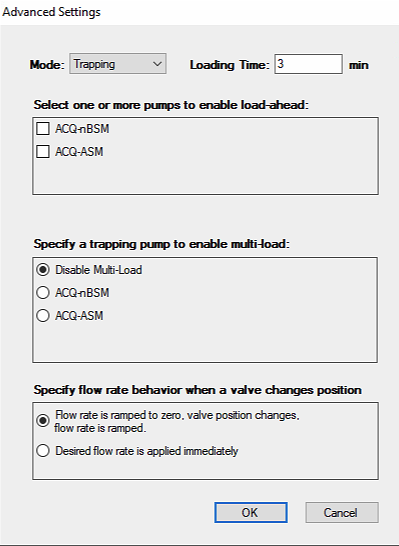

4. In the **ACQ-ASM Auxiliary Solvent Manager** perspective for **Trapping**:

- Set the **Run Time** to 130 min
- In the **Pump A Flow Table:**
  - Set the **Flow (µL/min)** to 10.000
  - Set the **Pressure Limits High** to 5,500 psi.
- In the **Pump B Flow Table** set the **Flow (uL/min)** to 0.000

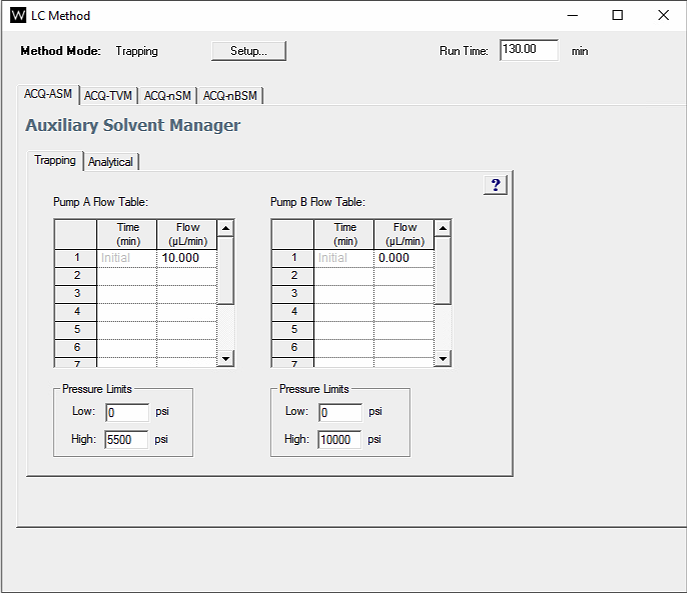

4. In the **ACQ-ASM Auxiliary Solvent Manager** perspective for **Analytical**:

- In the **Pump A Flow Table:**
  - Set the **Flow (µL/min)** to 1.000
  - Set the **Pressure Limits High** to 5,500 psi.
- In the **Pump B Flow Table:**
  - Set the **Solvent** to Acetonitrile
  - Set the **Flow (uL/min)** to 0.000

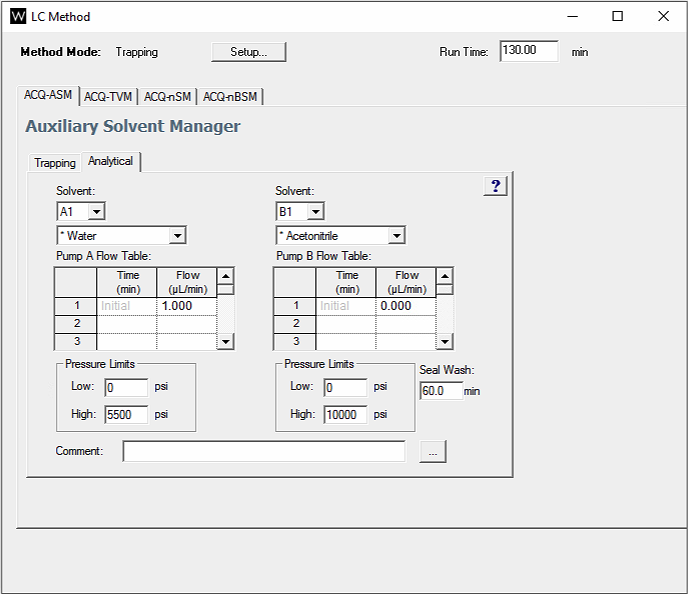

5. Navigate to the **ACQ-TVM Trap Valve Manager** perspective for **General**:

- Set the **Valve Positions** for the **Analytical** and **Trapping** configuration

*Note: Here the left valve is set to Position 1 for Analytical flow and Position 2 for Trapping*

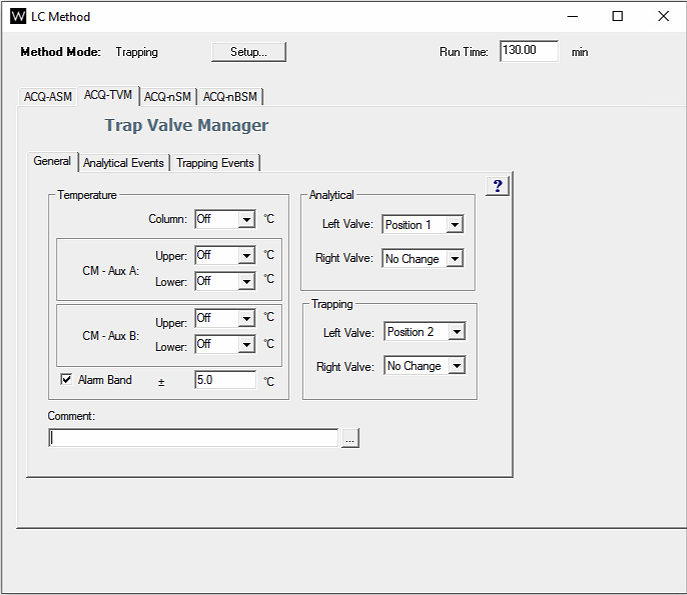

6. Navigate to the **ACQ-nSM µSample Manager - FL** perspective for **General**:

- Set the **Wash Solvents** and **Volumes**
  - **Weak**: 0.1% Formic Acid (FA) in Water, 600 µL
  - **Strong**: 0.1% FA in Acetonitrile (ACN), 200 µL
- Set the **Temperature Control** for the Sample to 10 °C

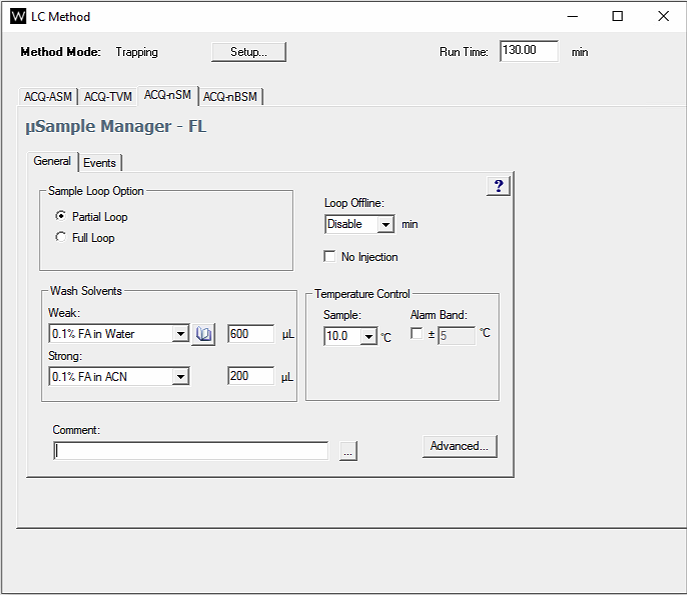

7. Navigate to the **ACQ-nBSM Binary Solvent Manager** perspective for **Trapping**:

- Set the **High Pressure Limit** to 6,000 psi
- Set the **Gradient**:
  - **Flow (µL/min)** to 5.000
  - **%A** to 95.0
  - **%B** to 5.0

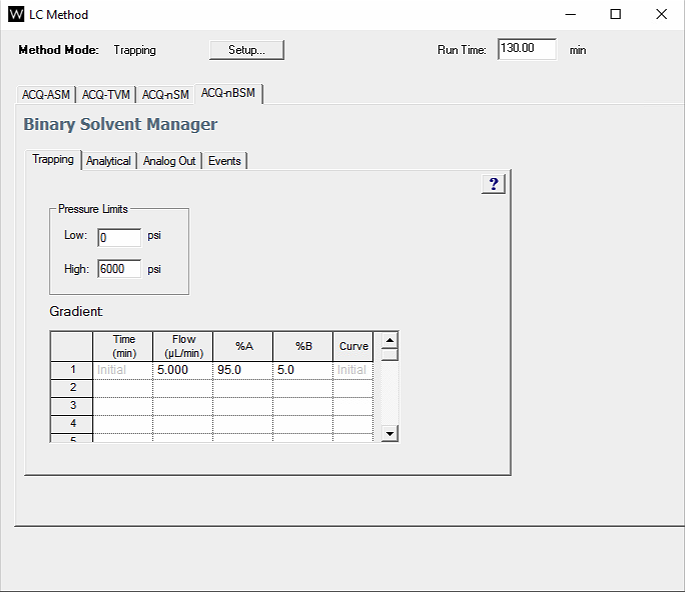

8. Navigate to the **ACQ-nBSM Binary Solvent Manager** perspective for **Analytical**:

- Set the **High Pressure Limit** to 6,000 psi
- Set the **Seal Wash** to 5.0 min
- Set the **Gradient** using the gradient table in Supplementary Table S1

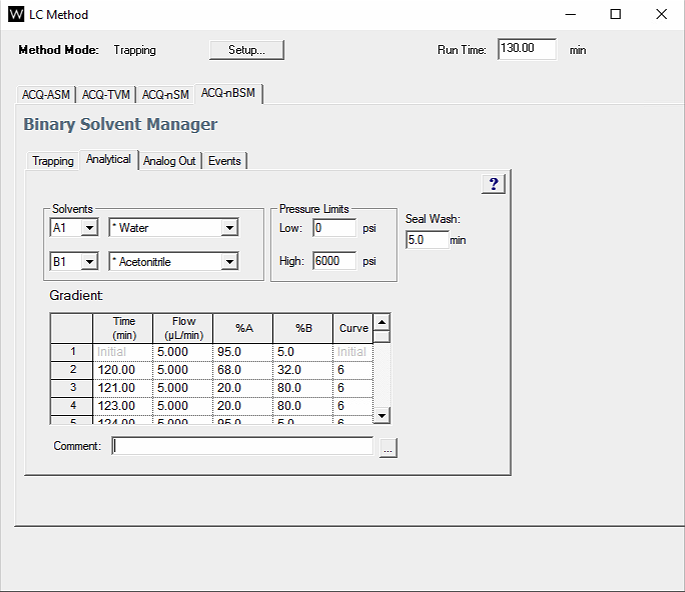

9. Save the LC Method by clicking the Floppy Disk Drive dropdown in the **LC Method** perspective and selecting “Save As”

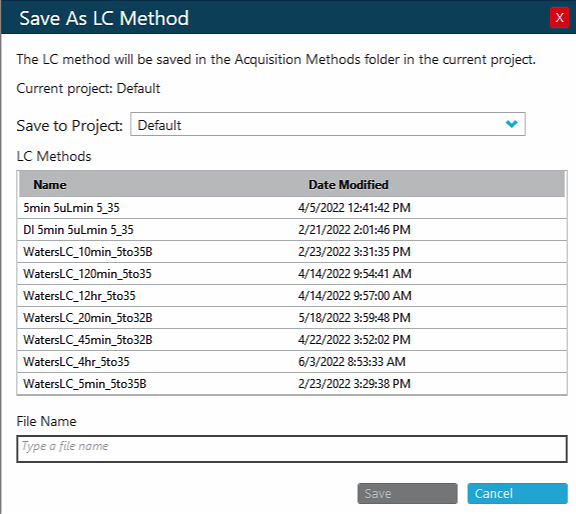

10. Name the method and click **Save**.

### **Chapter 5. Checking ZenoTOF 7600 Tuning Parameters in SCIEX OS**

1. Navigate to the **MS Tune** perspective.

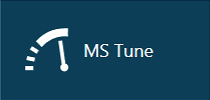

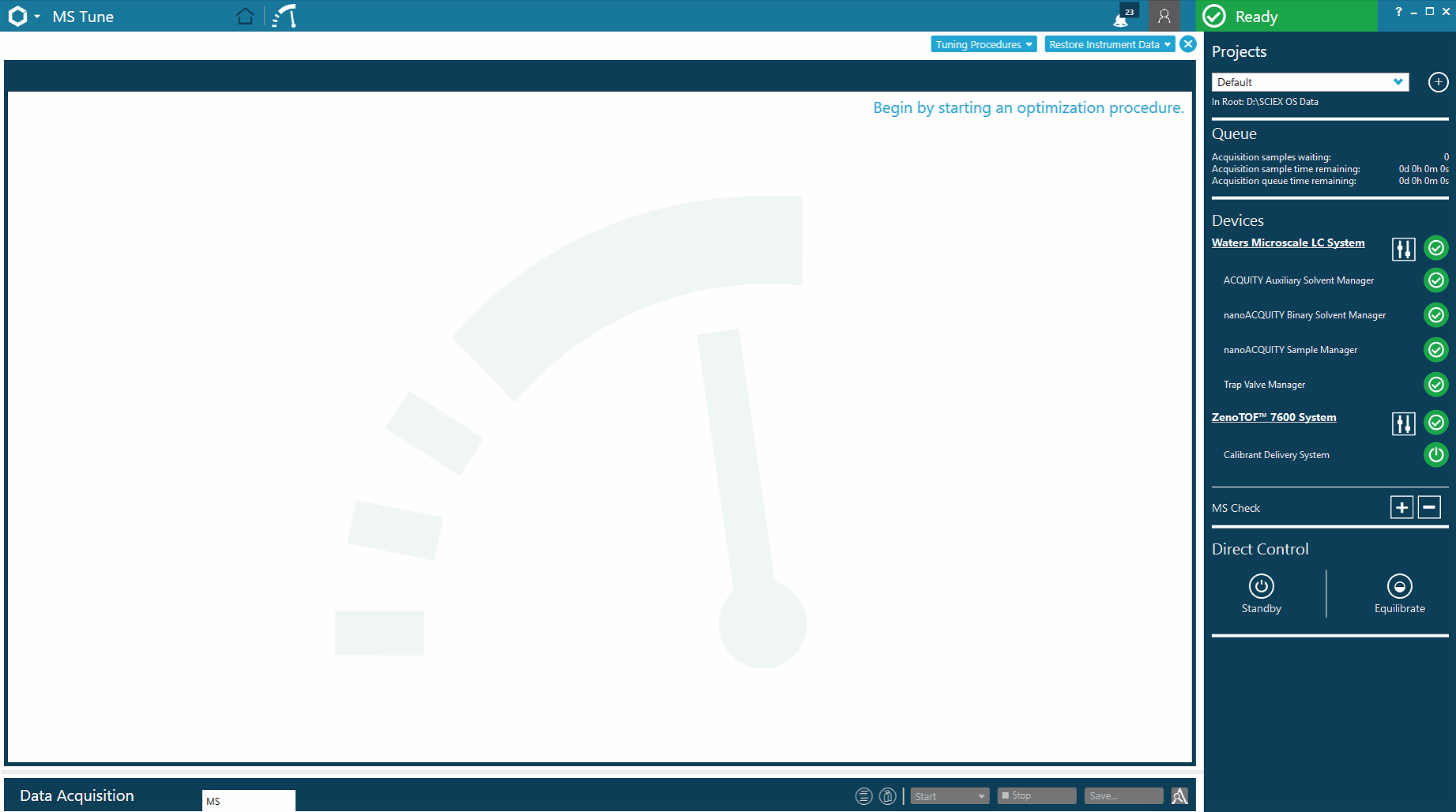

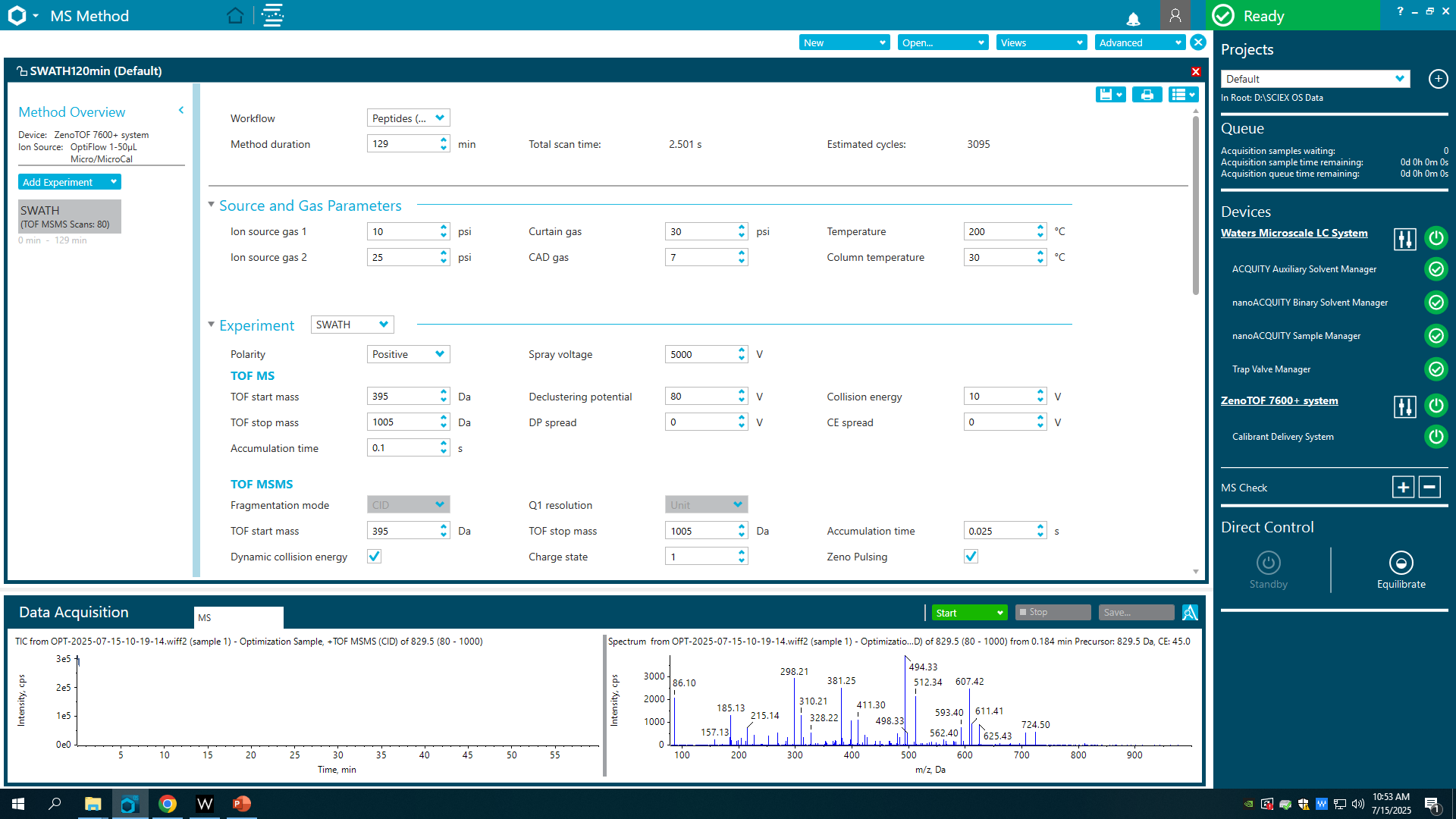

2. Select **Positive Quick Status Check** in the **Tuning Procedures** dropdown.

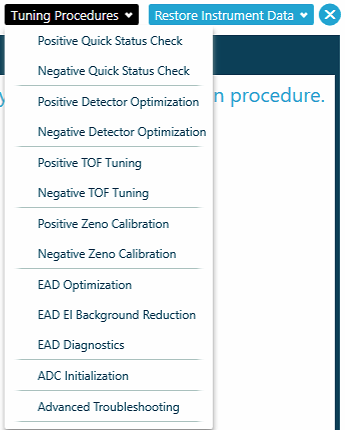

3. Ensure that the **CDS** containing the X500 ESI Calibration Solution is running and click **Next**.

*Note: The system will progress through the procedure automatically.*

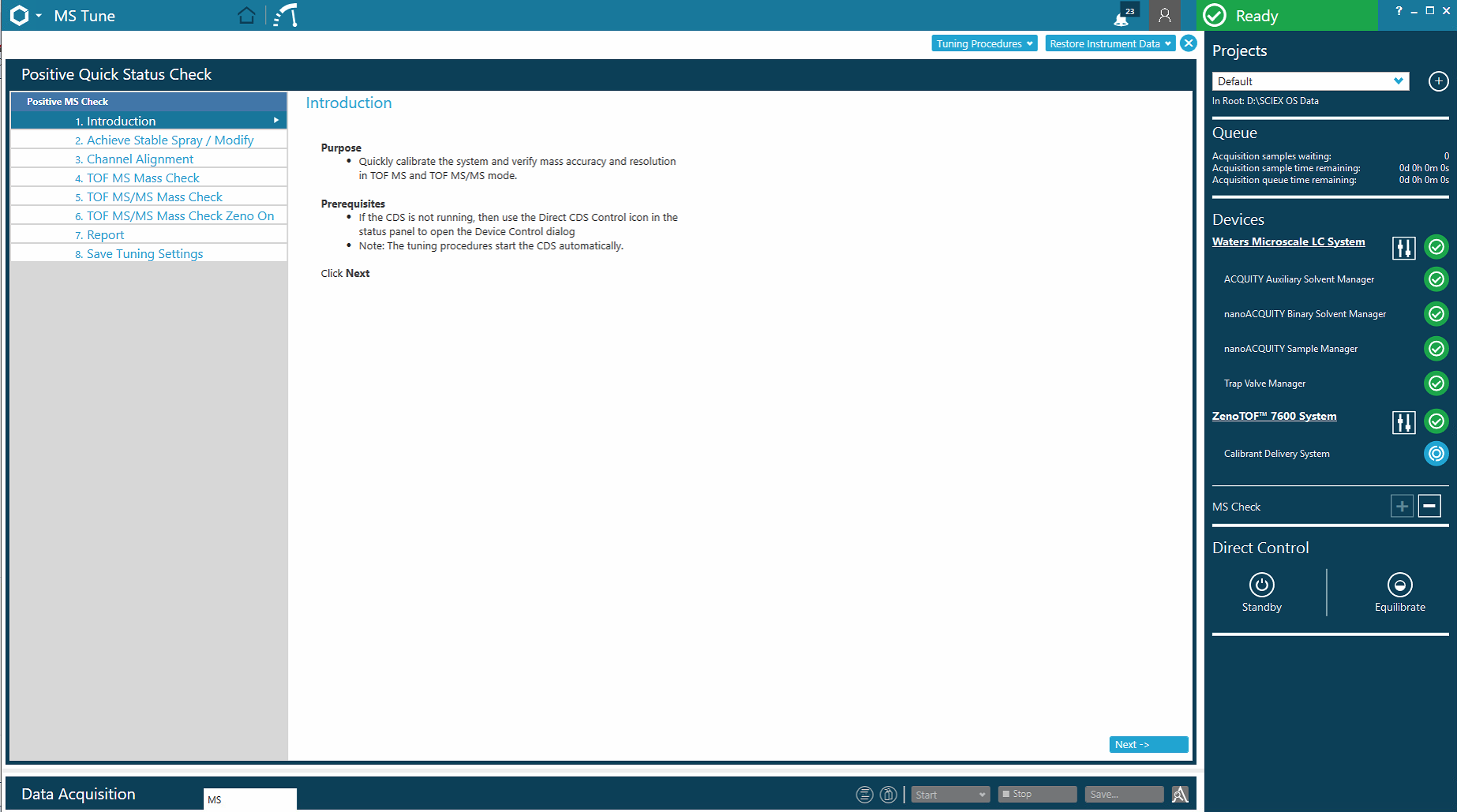

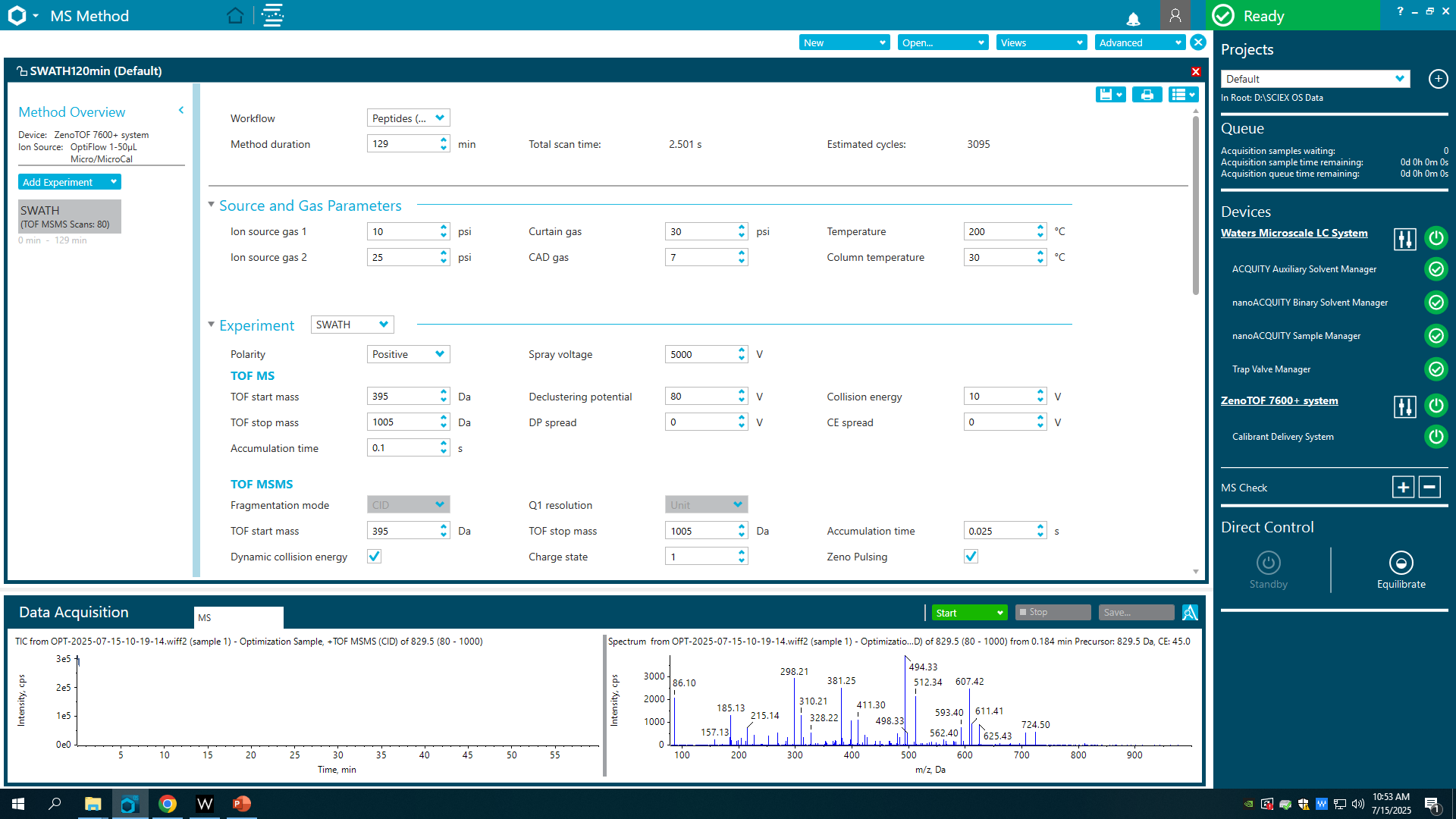

4. Assess the **Report** after the system achieves stable spray, aligns the channels, does mass checks.

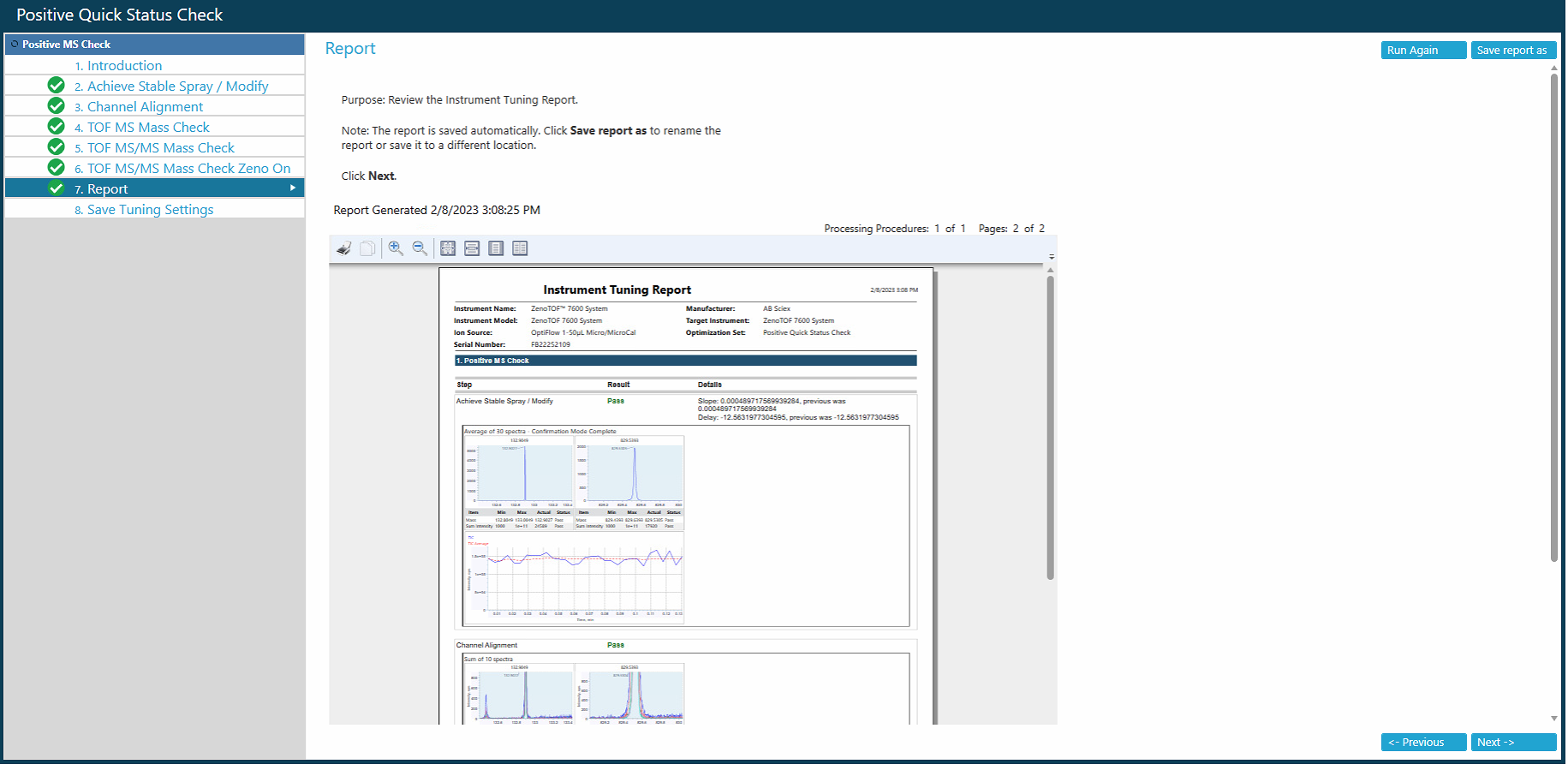

5. Click **Next.**

6. Save the Tune settings by clicking **Save Settings**.

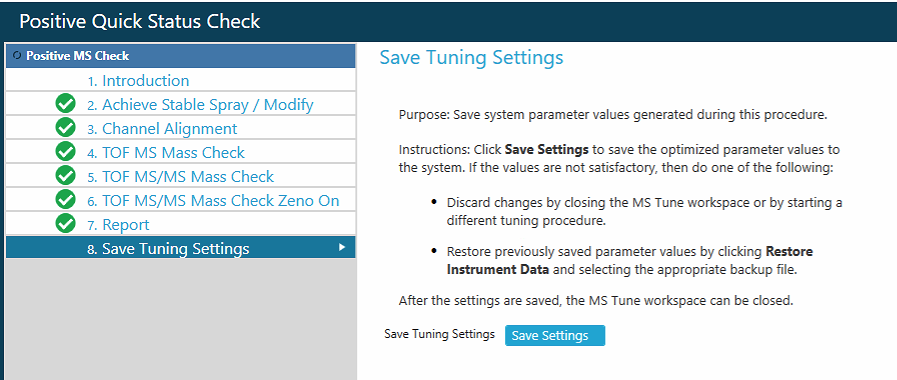

### **Chapter 6. Sample Batch Setup and Data Acquisition in SCIEX OS**

1. Open the **Batch** perspective from the SCIEX OS homepage.

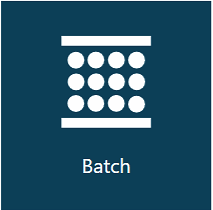

2. The **Batch** perspective has a fillable table. You can create a new batch or open one that was saved previously.

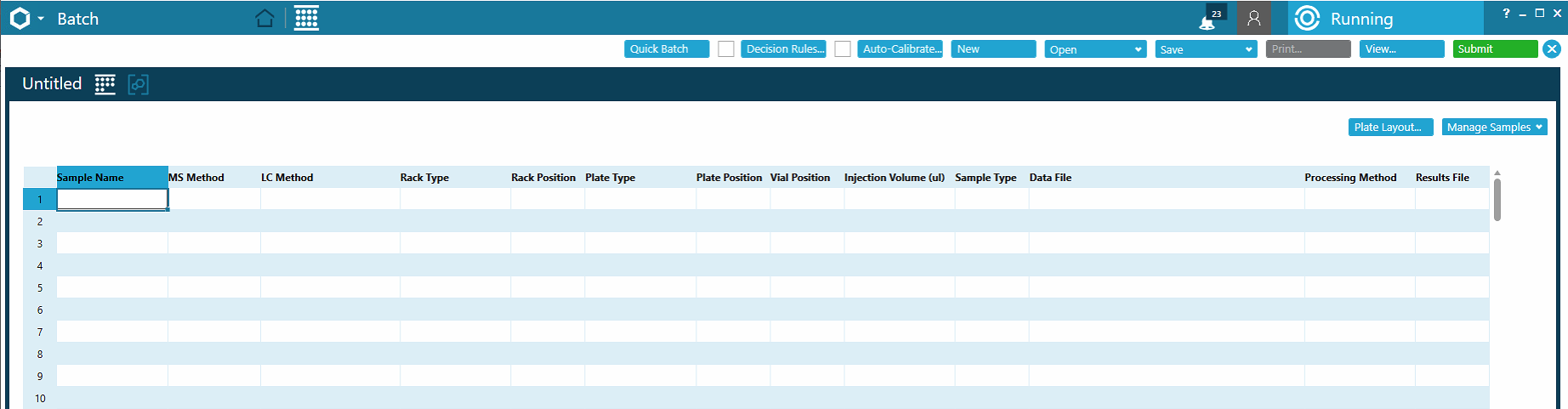

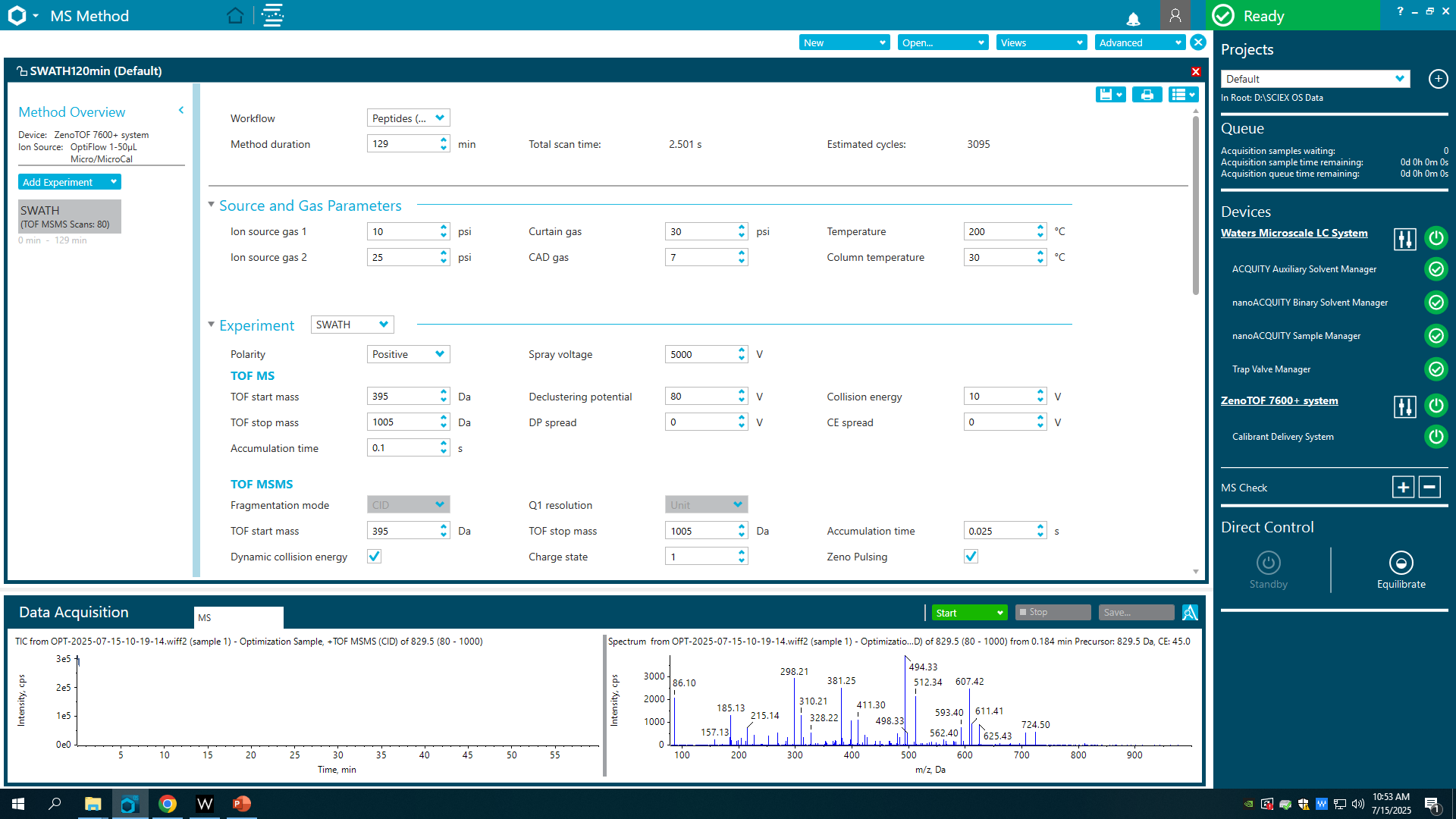

3. Fill in the **Batch** table:

- Input the **Sample Name** for each sample, blank, or quality control acquisition
- Set the **MS Method** to the desired method
- Set the **LC Method** to the desired method
- Set the **Rack Type** to “Sample Manager”
- Set the **Rack Position** to the appropriate tray
- Set the **Plate Type** to “ANSI-48Vial2mLHolder”
- Set the **Plate Position** to the appropriate tray
- Set the **Vial Position** to the appropriate position
- Set the **Injection Volume (µL)** to the desired injection amount
- Set the **Sample Type**
- Name the **Data File**
- Leave the **Processing Method** and **Results File** blank.

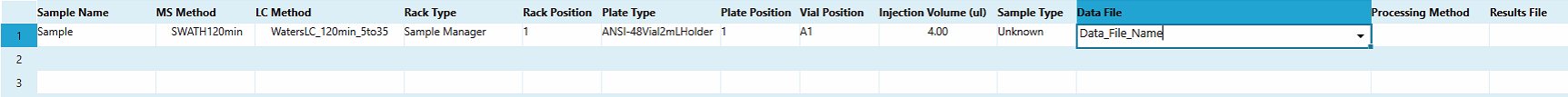

5. Enable the **Auto-Calibrate.**

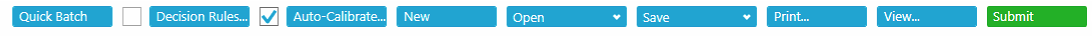

6. Enter the **Auto-Calibrate** perspective by clicking on the Icon:

- Set the **Ion reference table** to “X500 ESI Positive Calibration Solution
- Set the **Calibrate every** to the appropriate number of samples (here we use 5 samples)
- Set the **Calibrant delivery** to “CDS”
- Set the **CDS** channel to the appropriate bottle of calibrant (here we use channel 1).

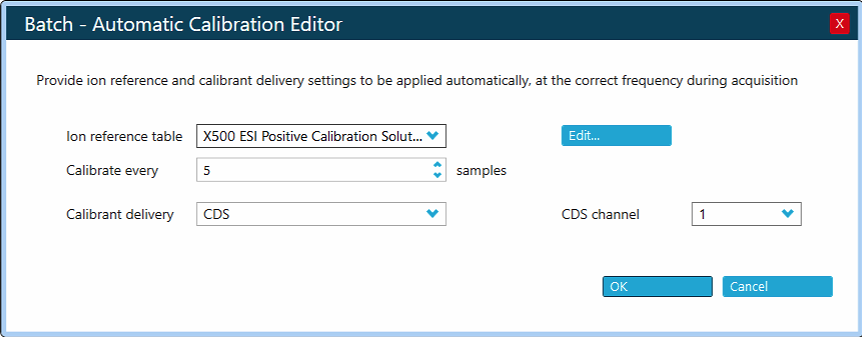

7. Click **OK**.

8. Save the batch by clicking the **Save** icon and **Save As**.

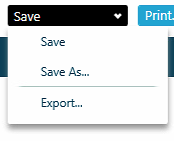

9. Input an appropriate name for the Batch and click **Save**.

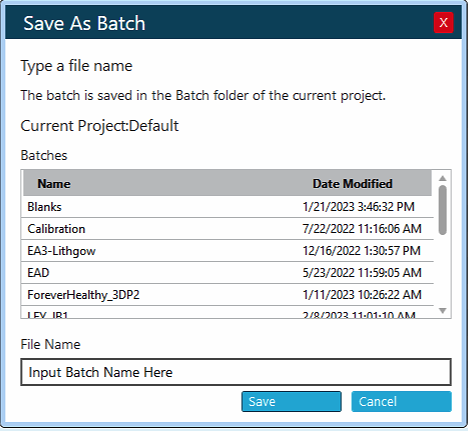

10. Click **Submit**.

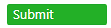

11. The system should change from **Ready** to **Running** in the upper left corner.

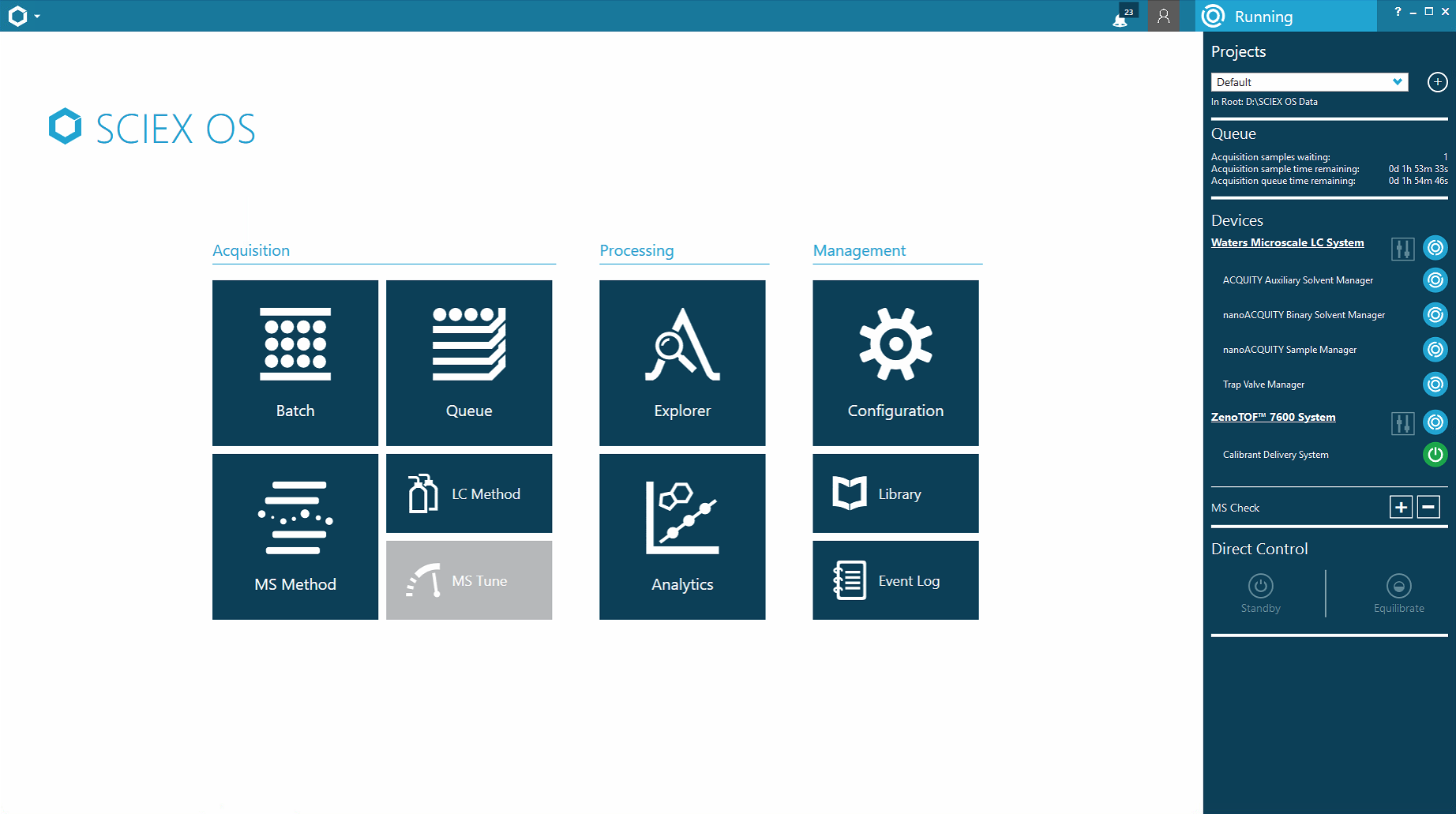

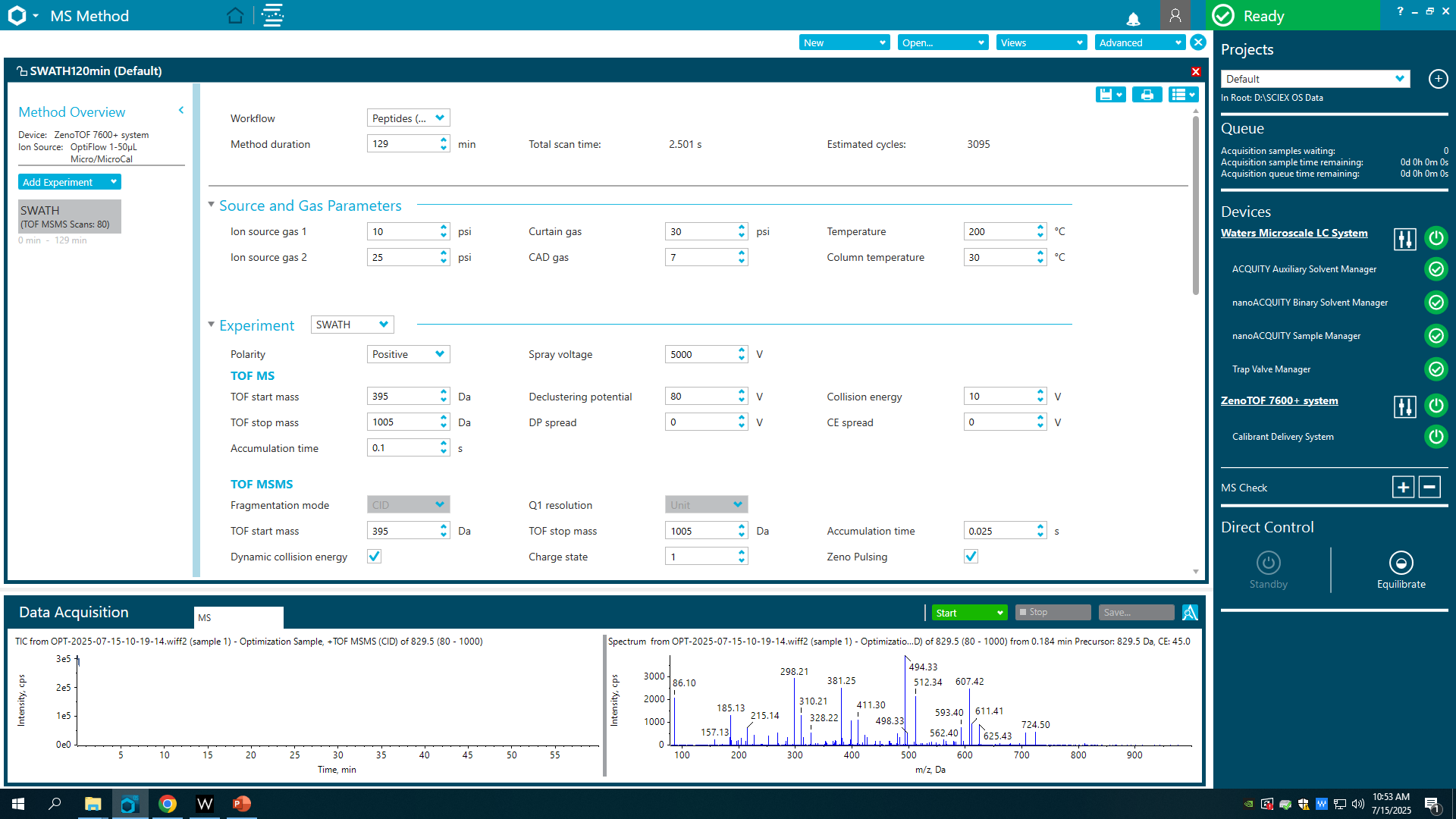

12. Open the **Queue** perspective.

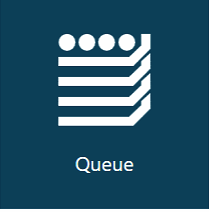

13. Monitor Batch acquisition progress in the **Queue** perspective.

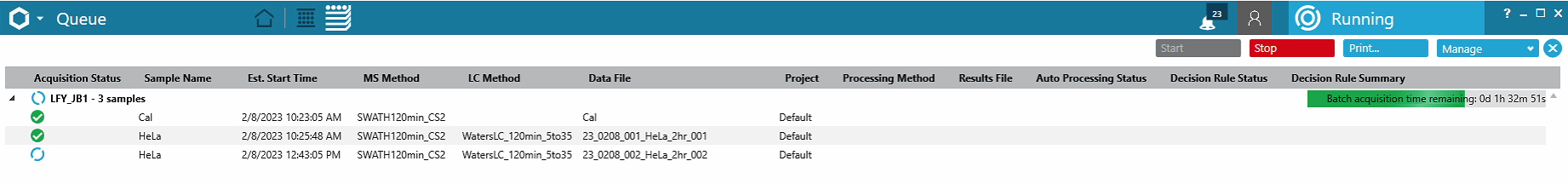

14. Click the **Data Acquisition** bar at the bottom of **SCIEX OS** to see the acquisition in real time.

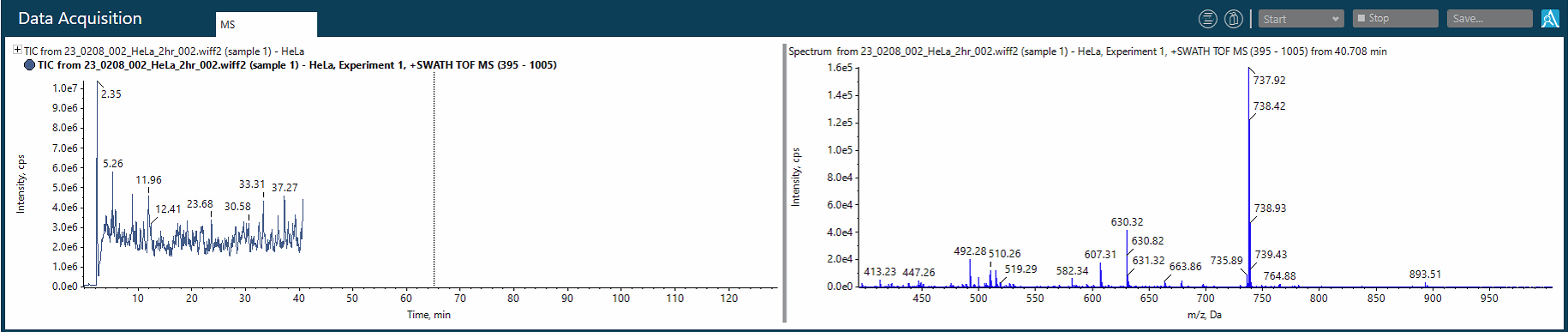

15. Double click a previously acquired sample in the **Queue** to view the data in the **Explorer** perspective.

- Top graph shows the **Total Ion Current**
- Open the bottom graph by clicking **Show, Base Peak Chromatogram,** and **Ok**.

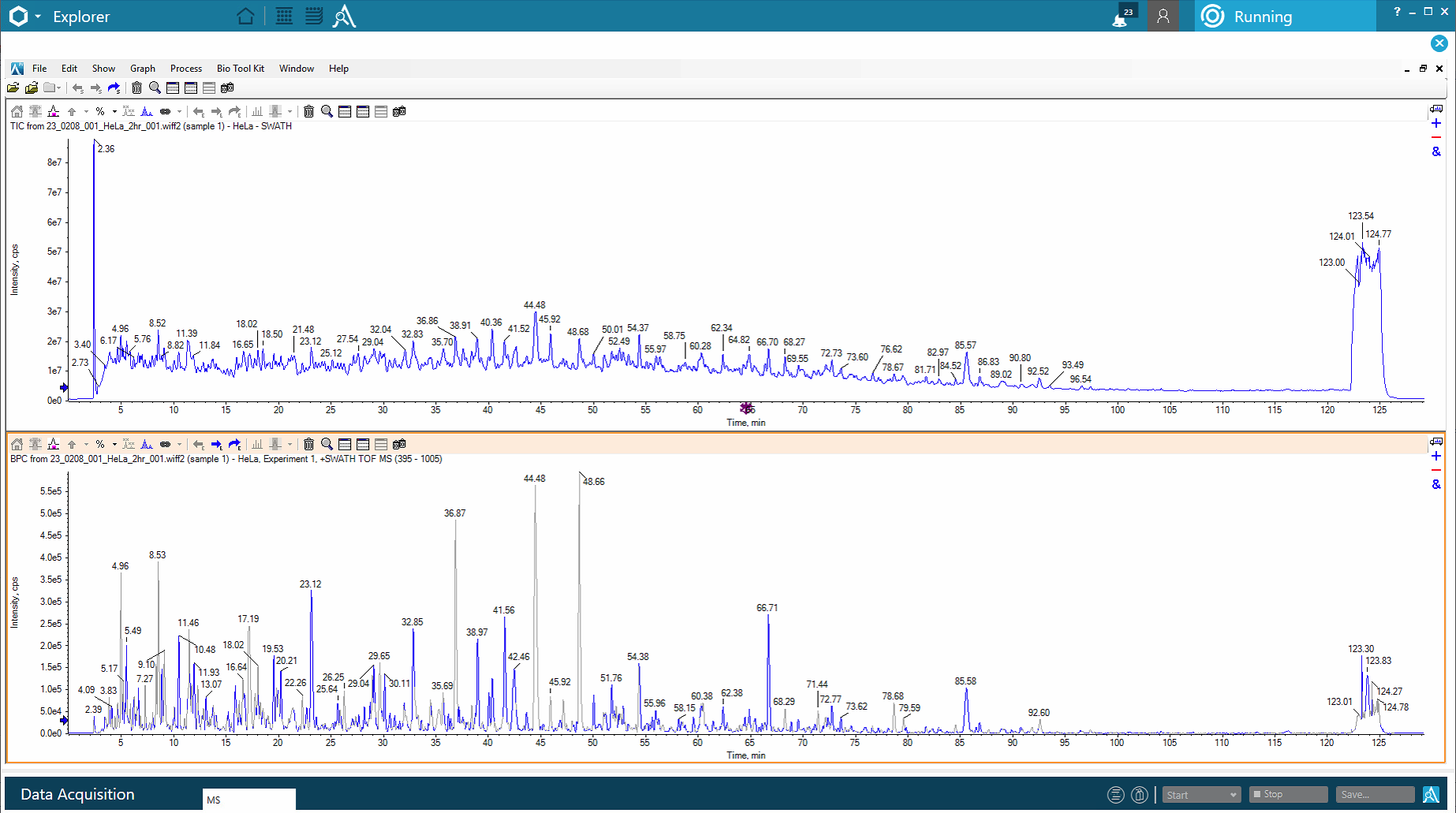

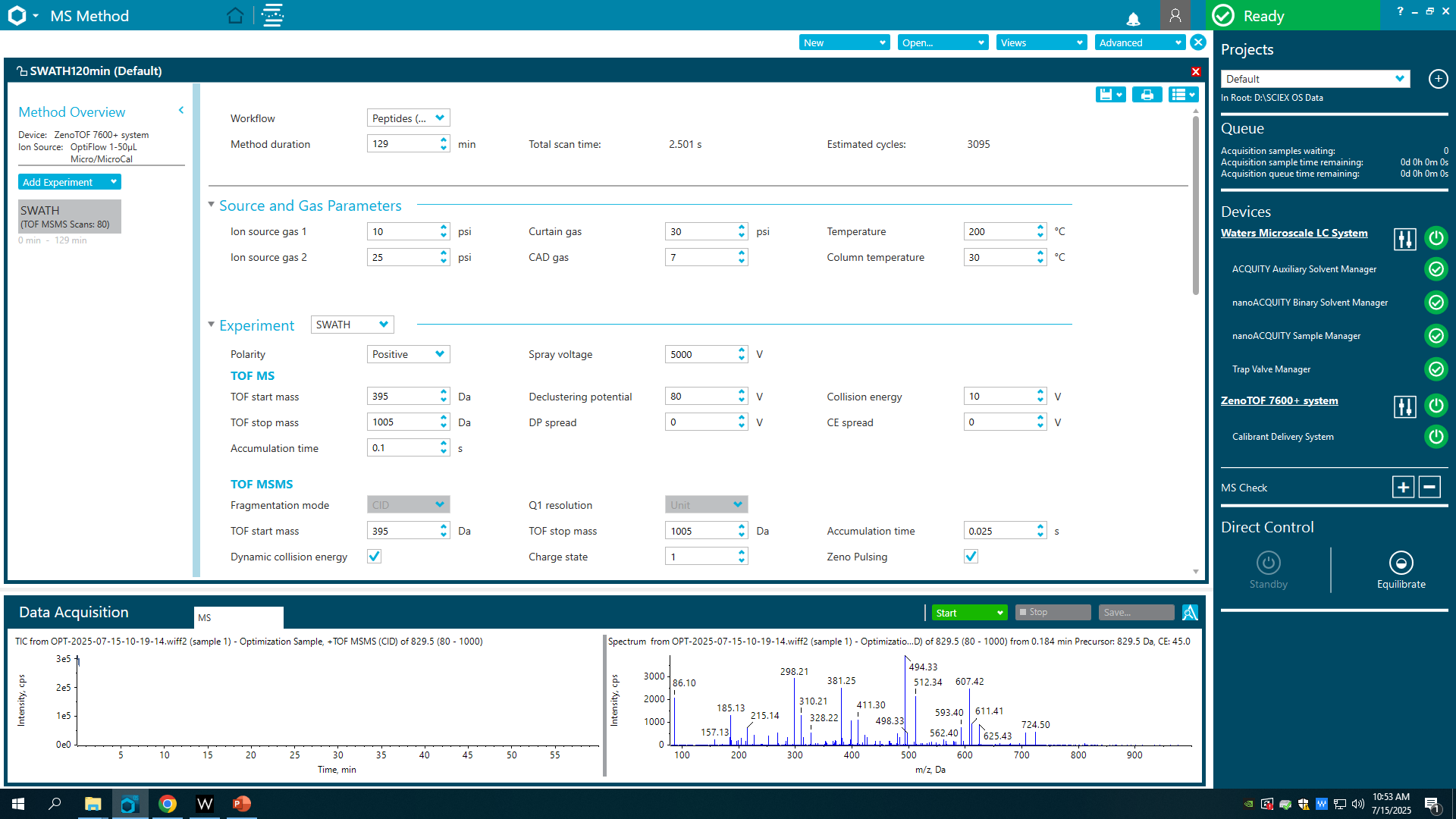

### **Chapter 7. Data Pre-Processing (Optional Step)**

Once the DIA data is acquired, there are several data processing pipelines available to process the data. As an example, a DIA data set was acquired using the above method, then pre-processed using the HTRMS Converter program before directDIA analysis using Spectronaut (Biognosys, Schlieren, Switzerland). The HTRMS file conversion is optional and decreases the time to search the data using Spectronaut by converting vendor specific file formats into a **H**igh **T**ime **R**esolution **M**ass **S**pectrometry (HTRMS) file format. The optional file conversion step is optimal for large sample sizes.

1. Install the HTRMS Convertor program from Biognosys onto the MS computer.

*Note: This requires a restart.*

2. Open the **HTRMS Convertor program** to set the program up to convert acquired data-independent acquisition files to a *.htrms peaks list in a monitored folder.

3. Select **Add Folder**.

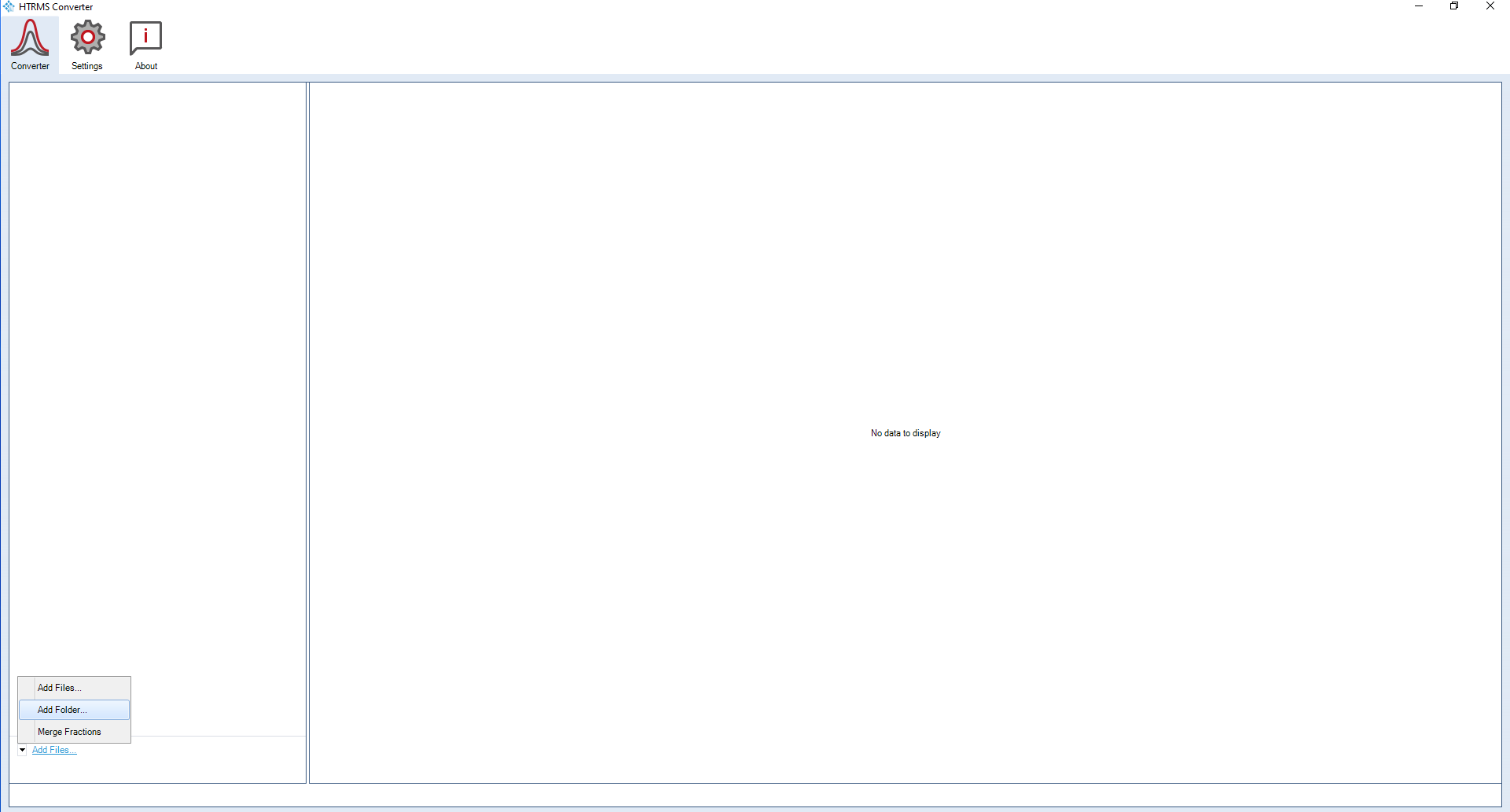

4. Select the **Source** and **Destination Directories**.

5. Ensure that **Monitor Folder** is enabled.

6. Click **Ok**.

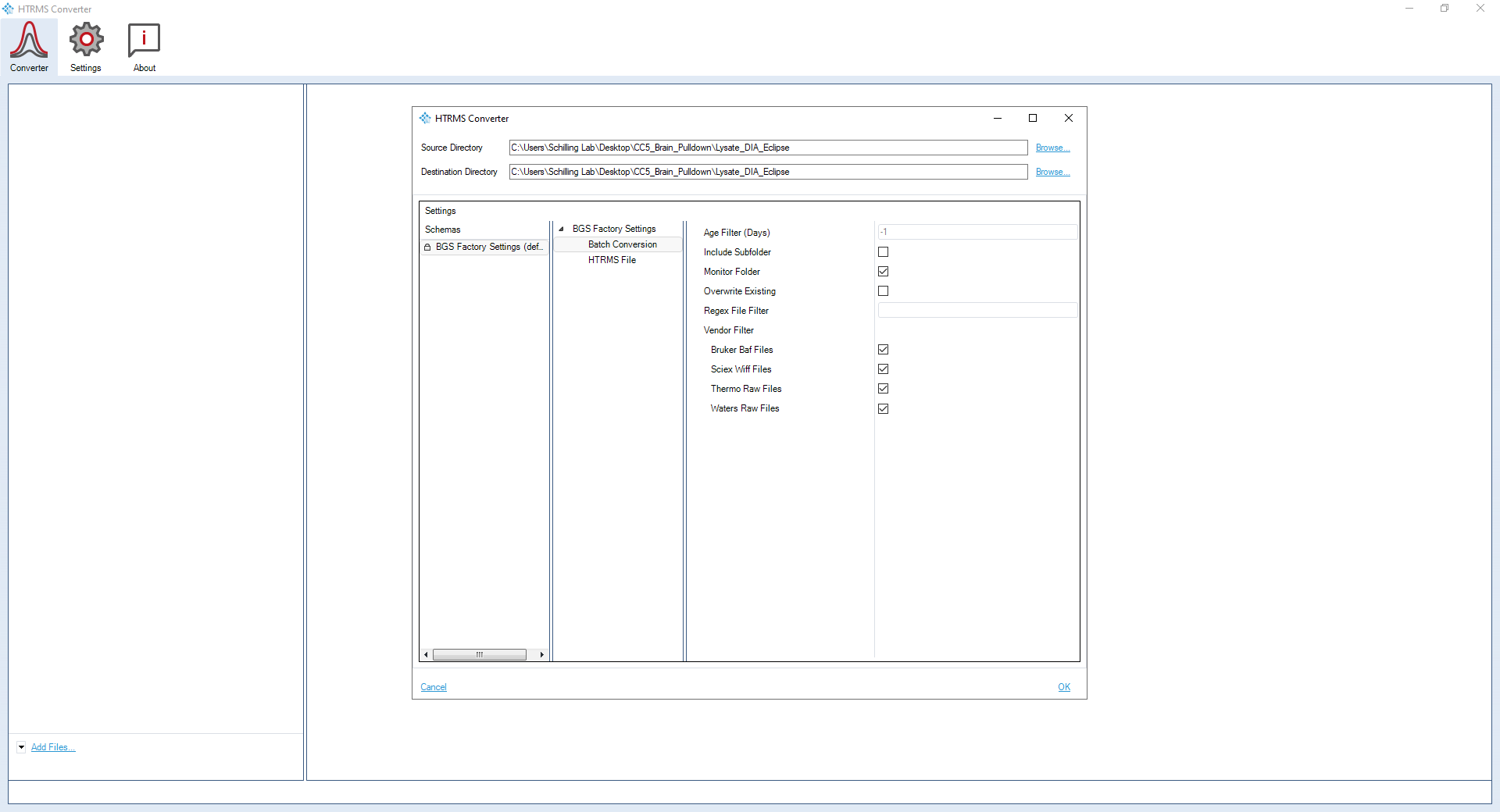

### **Chapter 8. Data Processing using Spectronaut**

Spectronaut data processing using directDIA analysis searches each sample acquisition to form a spectral library, then searches each sample acquisition against the library to identify precursors, peptides, proteins, and protein groups.

1. Open **Spectronaut.**

*Note: This tutorial is based on Spectronaut version 17.1.221229.55965.*

2. Select the **Pipeline** tab:

- click **Set up a directDIA analysis from File**
- navigate to the experimental vendor specific files or *.htrms files and select all files pertinent to the experiment
- Click **Open**.

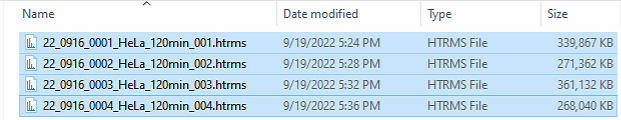

3. Name the experiment with a unique identifier.

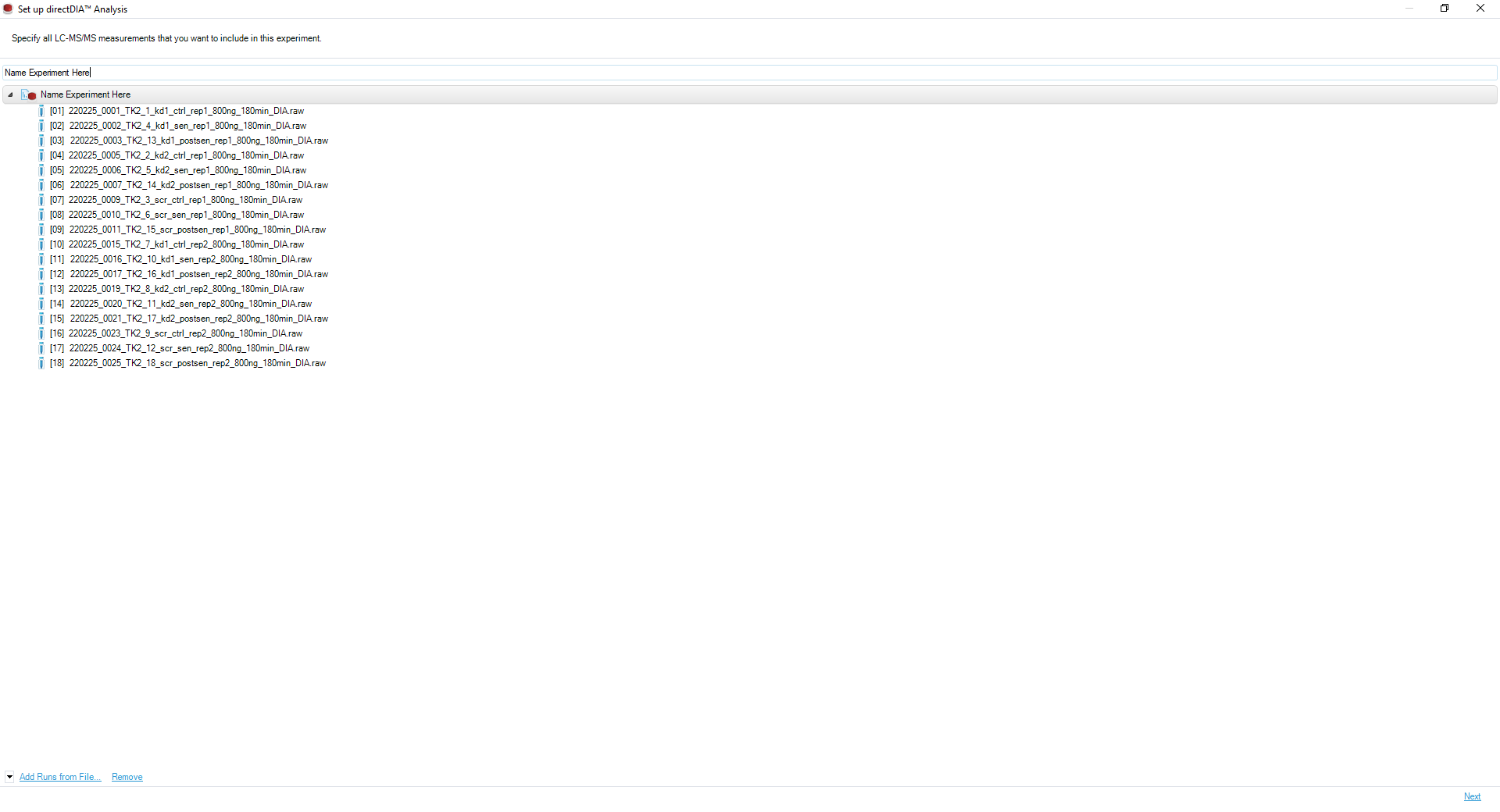

4. Click **Next**.

5. Select the appropriate organism **protein database**.

*Note: In this study, the FASTA file “uniprot-human proteome_reviewed” is used.*

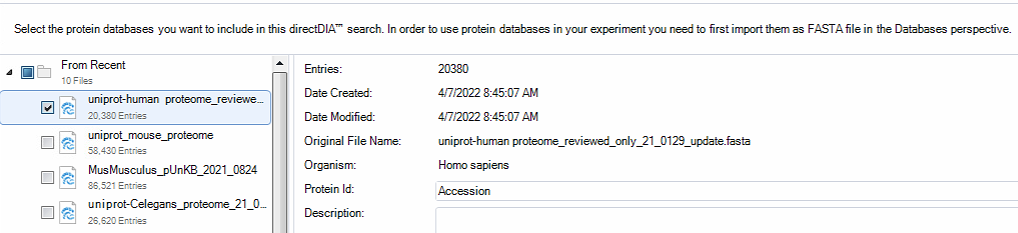

6. Click **Next**.

**Chapter 8.1. Protein Lysate: directDIA Search Settings**

Create a **new search and extraction settings schema** for directDIA analysis of a digested protein lysate without specific post-translational modifications. Defaults should be preserved unless specifically mentioned in the text below.

1. Navigate to the **Pulsar Search Peptides** tab:

- the *Enzymes/Cleavage Rules* should be set to “Trypsin/P”
- set the *Digest Type* to “Specific”
- set the *Missed Cleavages* to “2”
-
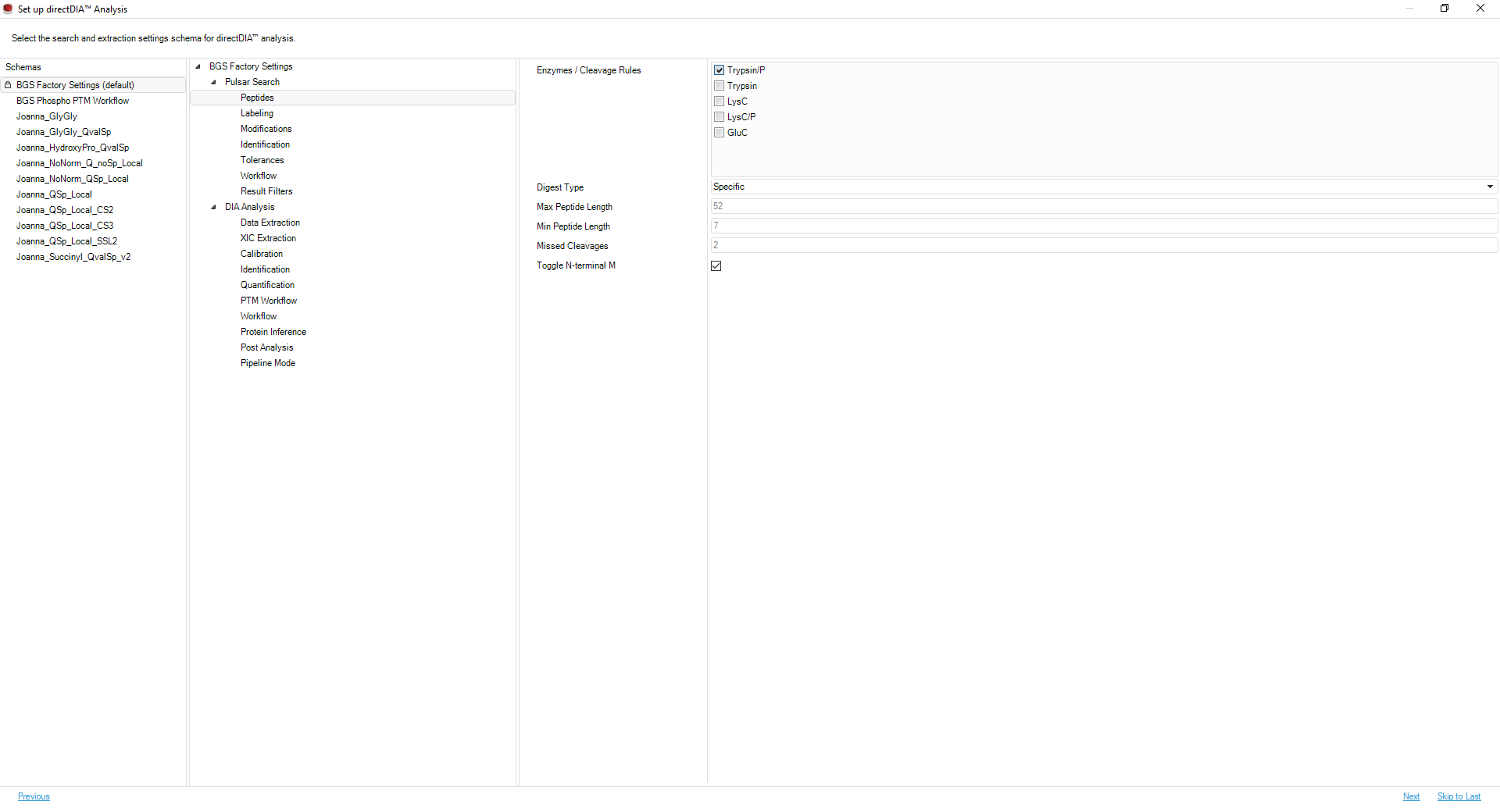
enable *Toggle N-terminal M.*

2. Select the **Pulsar Search Modifications** tab:

- set the *Max Variable Modifications* to “5”
- set the *Fixed Modifications* to “Carbamidomethyl (C)”
- set the *Variable Modifications* to “Acetyl (Protein *N*-Term)” and “Oxidation (M)”.

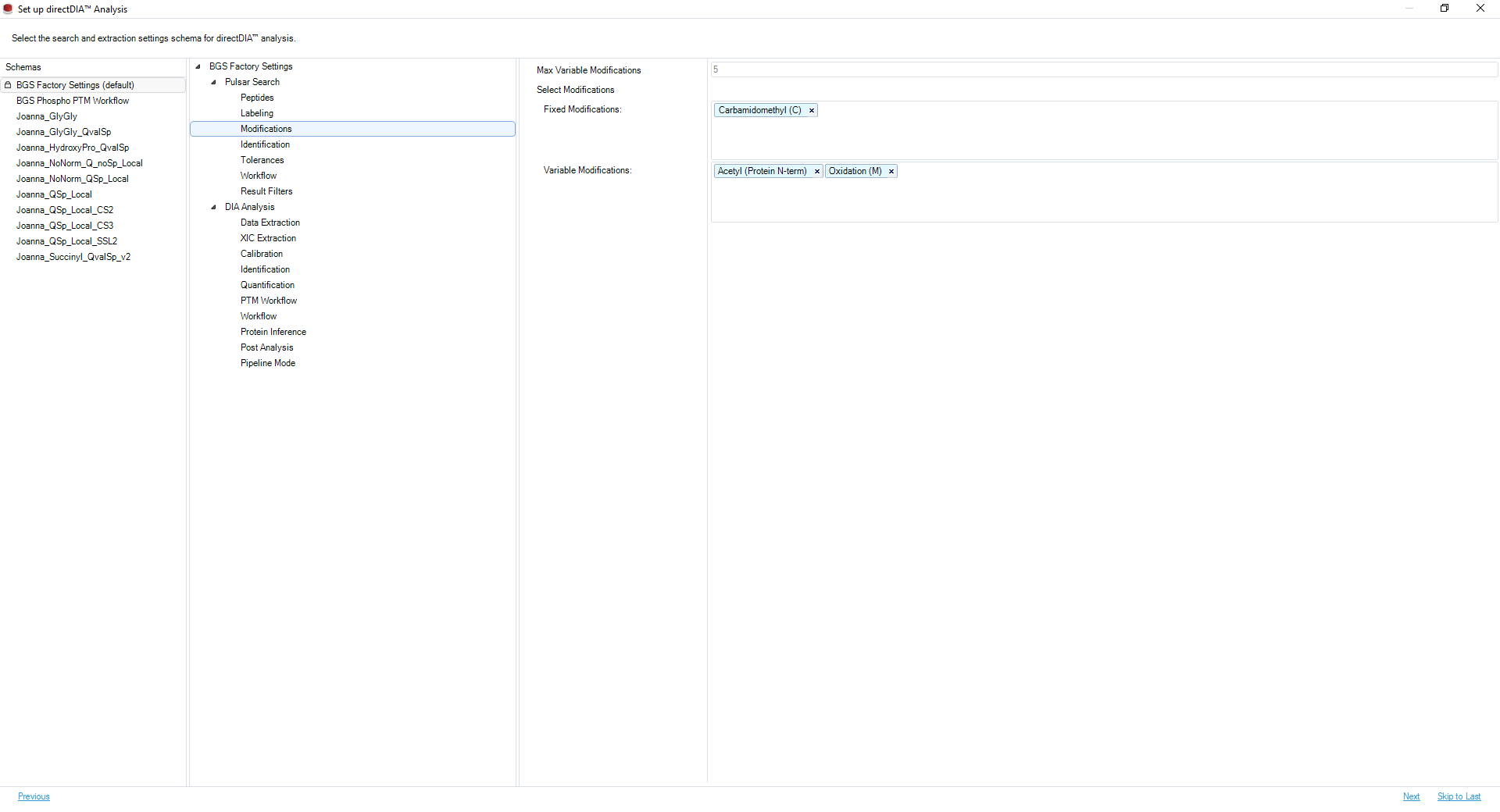

3. Navigate to the **DIA Analysis Identification** tab:

- set the *Precursor PEP Cutoff* to “1”
- set the *Precursor Qvalue Cutoff* to “0.01”
- set the *Protein Qvalue Cutoff (Experiment)* to “0.01”
- set the *Protein Qvalue Cutoff (Run)* to “0.05”
- enable *Exclude Single Hit Proteins*.

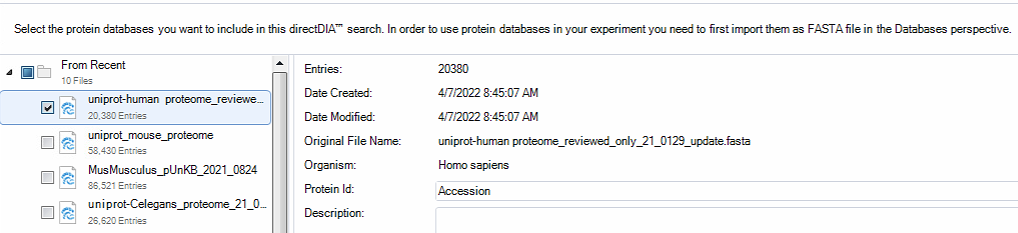

4. Click on the **DIA Analysis Quantification** tab:

- set the *Precursor Filtering* to “Identified (Qvalue)”
- set the *Imputing Strategy* to “None”
- set the *Normalization Strategy* to “Local Normalization”
- set the *Row Selection* to “Identified in at least 1 Run (Sparse)”.
- set the *Minor (Peptide) Grouping* to “by Stripped Sequence”
- set the *Major Group Quantity to “*Sum peptide quantity*”*
- set the *Major Group Max* to “7”
- set the *Major Group Min* to “1”
- set *Minor Group Quantity* to “Sum precursor quantity”
- set the *Minor Group Max* to “10”
- set the *Minor Group Min* to “1”

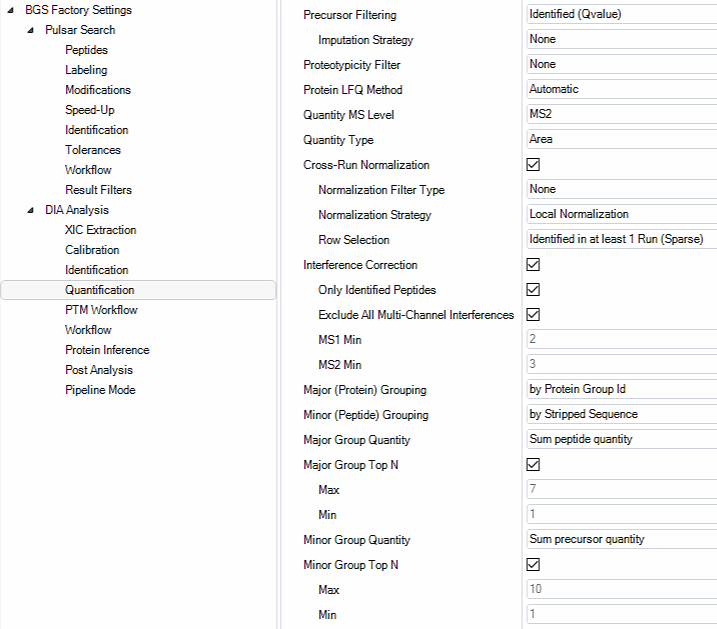

5. Select the **DIA Analysis Workflow** tab”

- set the *Profiling Strategy* to “iRT Profiling”.

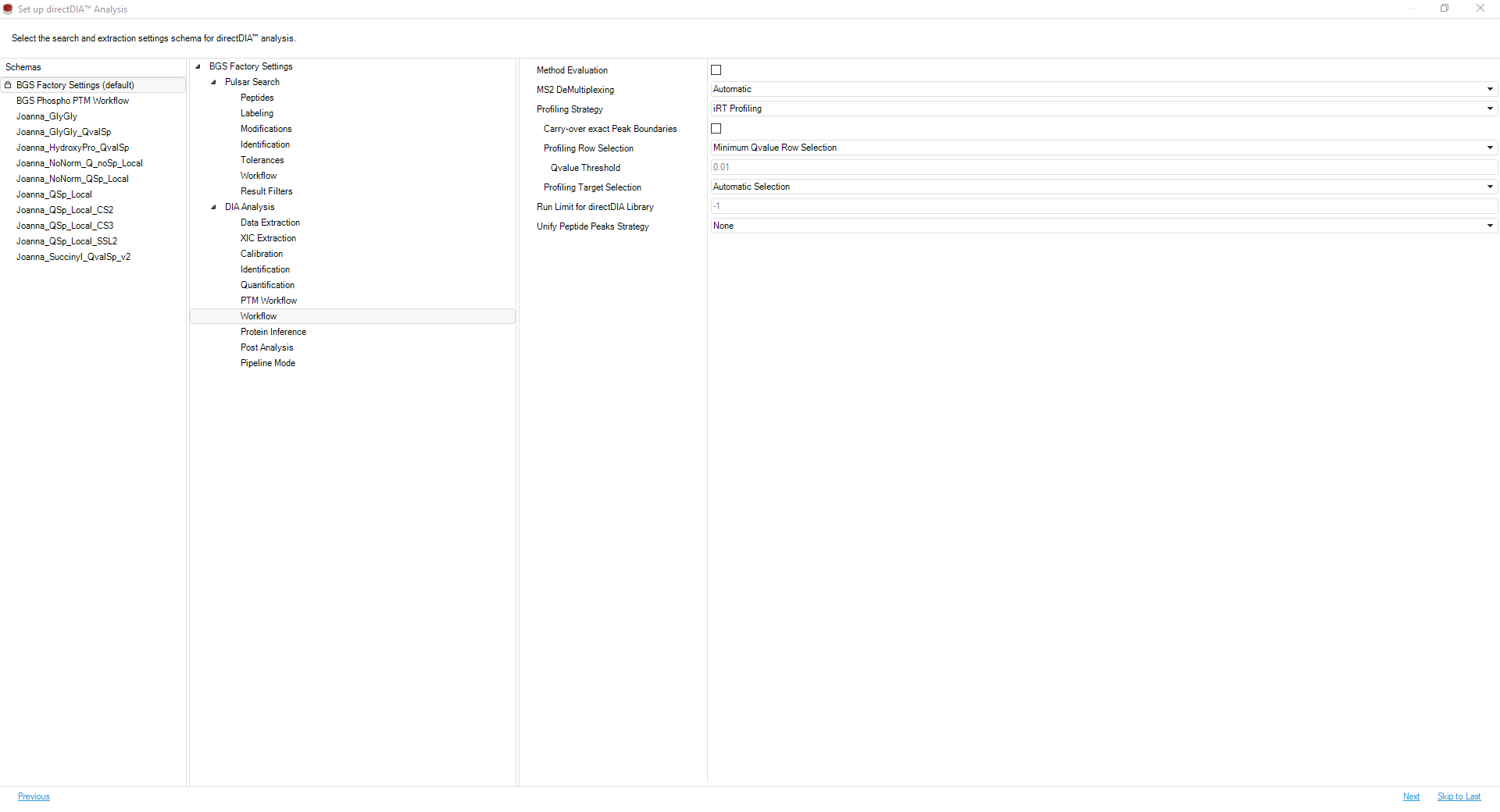

6. Navigate to the **DIA Analysis Post Analysis** tab:

- set the *Smallest Quantitative Unit* to “Precursor Ion (Quantification Settings)”
- set the *Differential Abundance Testing* to “Paired t-test”
- enable *Group-Wise Testing Correction*.

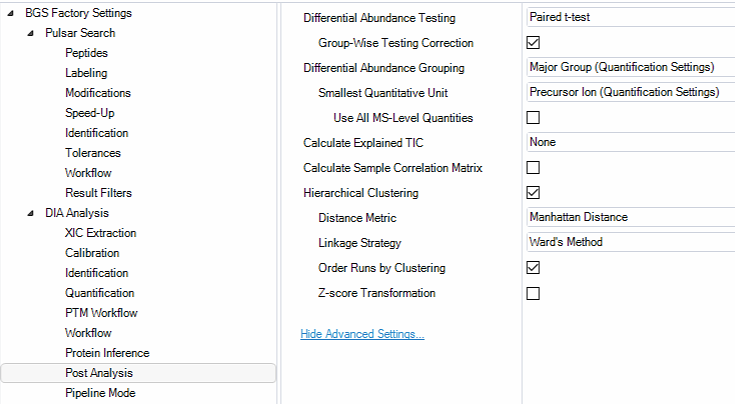

7. Click on the **DIA Analysis Pipeline Mode** tab:

- enable *Generate SNE File*
- select the appropriate *Report Schema* as described in **Chapter 4. Data Reporting***: “*Protein_Quant_Pivot” and “Peptide_Quant_Pivot”.

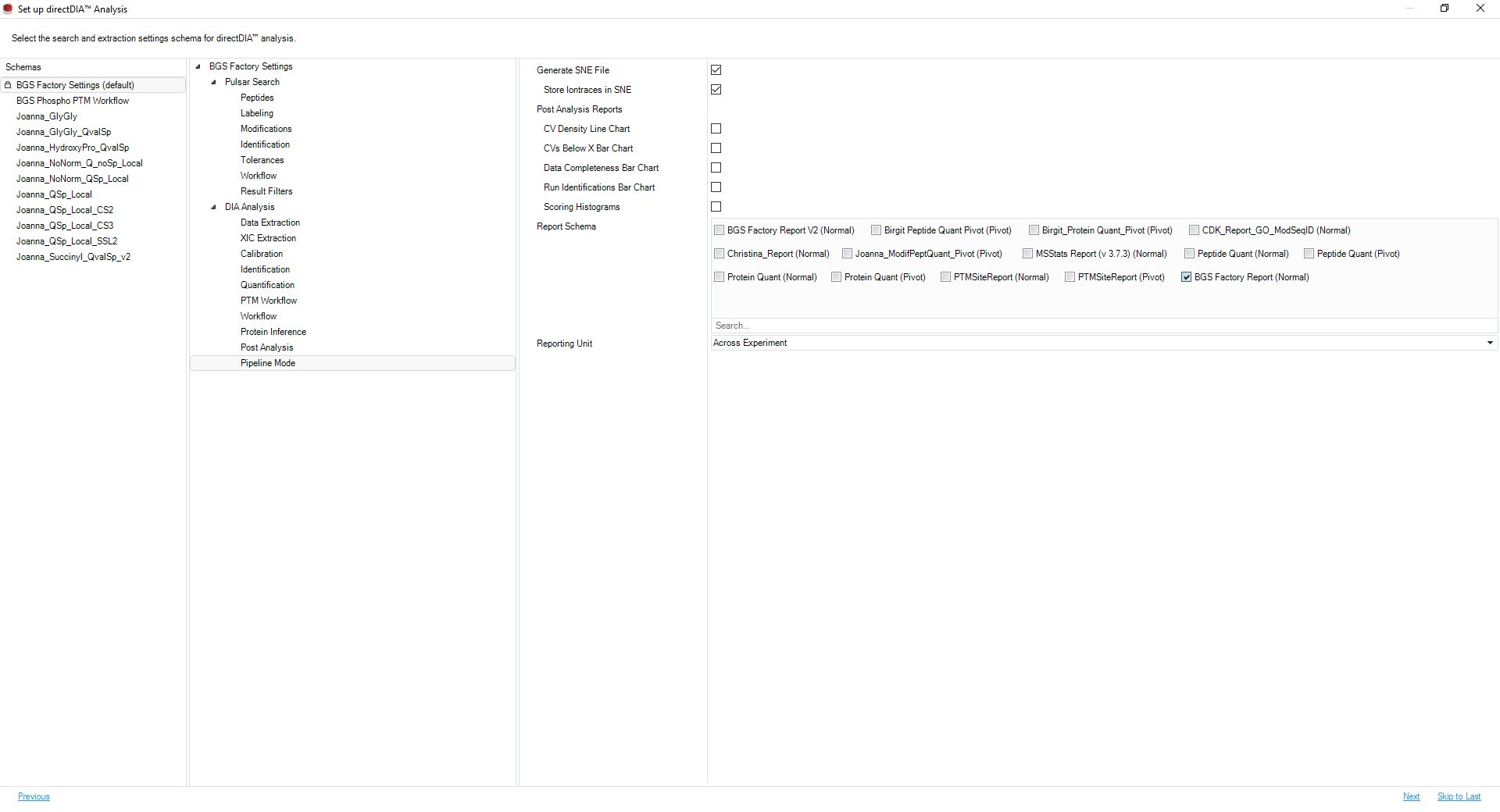

8. Click **Next**.

9. Go to **Chapter 3.3** to complete modifying search settings.

#### **Chapter 8.2. Comparison Setup, Gene Ontology Annotations, and Start Process**

1. Set the conditions for each experimental vendor specific file or *.htrms file included in the experiment.

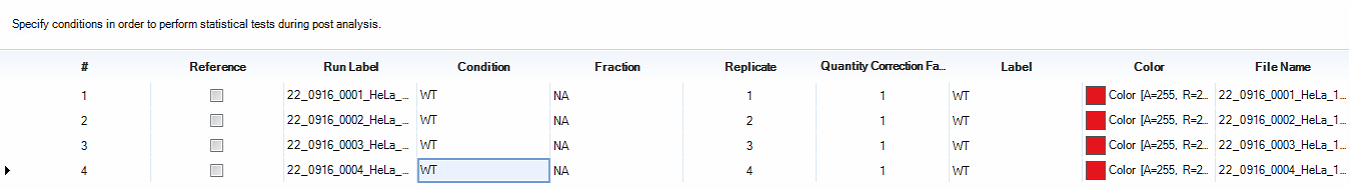

2. Click *Export Condition Setup*, name the file, and click **Save**.

3. Click **Next**.

4. Select the appropriate **Gene Ontology Annotations** for the experiment.

*Note: In this study, the file* “Homo Sapiens (GO Annotations Uniprot)” *is used.*

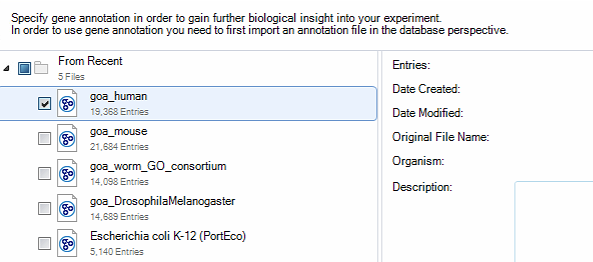

5. Click **Next**.

6. Click **Next** to advance past the Library Extension Runs page**.**

6. Click **Browse**:

- select the appropriate **Output Directory** to save the sne. file and export the reports, plots, and candidate protein files.

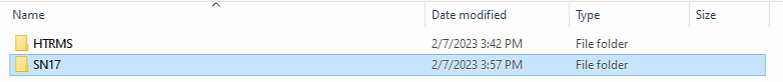

7. Click **Ok**.

8. Click **Finish** to queue the analysis.

9. Click **Run Pipeline** to start the analysis queue.

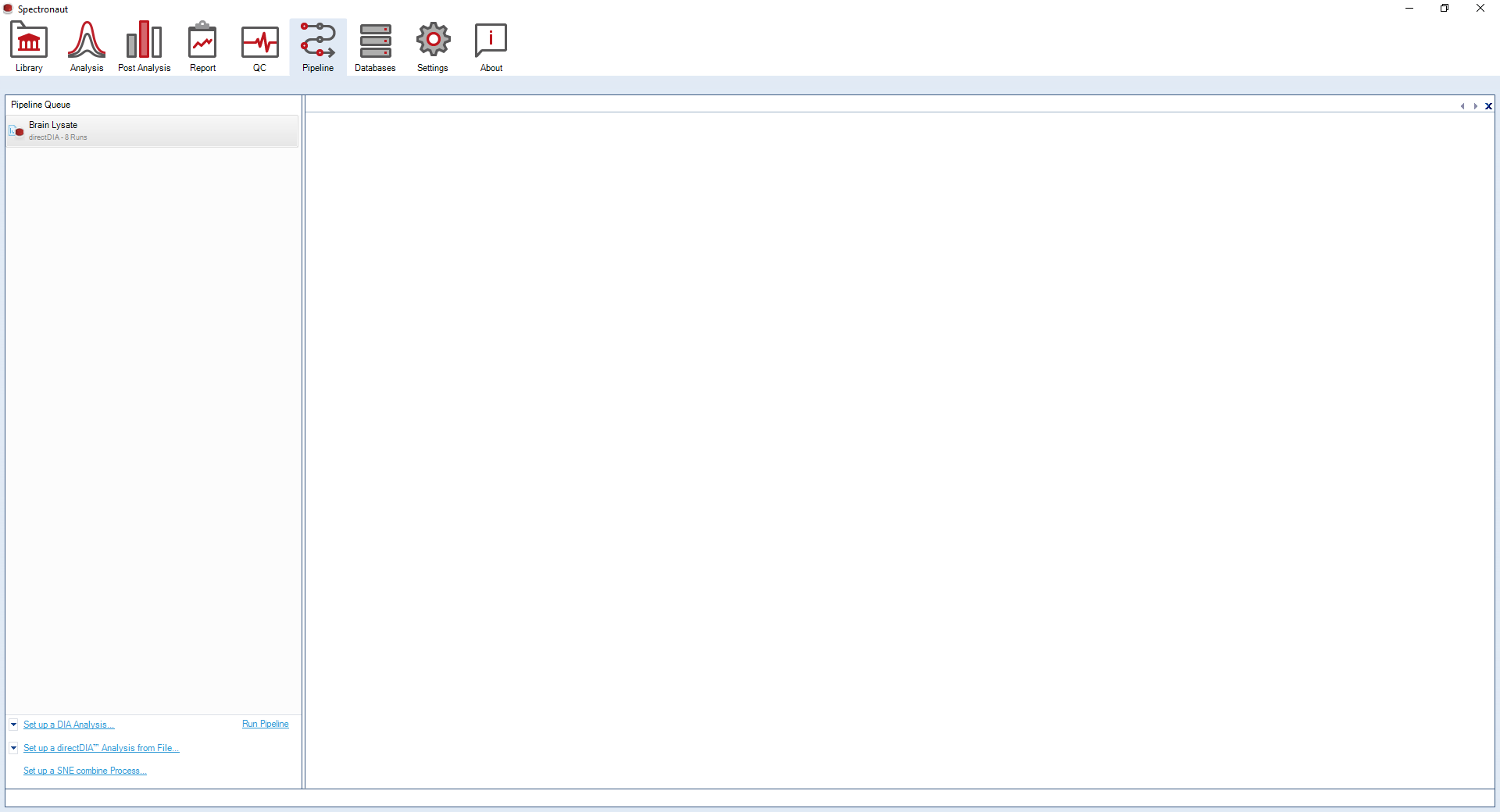

### **Chapter 9. Data Reporting**

#### **Chapter 9.1. Protein Quantification Report**

1. Navigate to the **Report** tab to set up a custom report.

2. Select the **BGS Factory Report** under the **Run Pivot Report** dropdown to create a custom Protein Quantification Pivot Report, “Protein_Quant_Pivot”.

3. Select the following **Row Labels**:

- PG.Qvalue
- PG.Genes
- PG.ProteinDescriptions
- PG.UniProtIDs
- PG.ProteinNames
- PG.CellularComponent
- PG.BiologicalProcess
- PG.MolecularFunction.

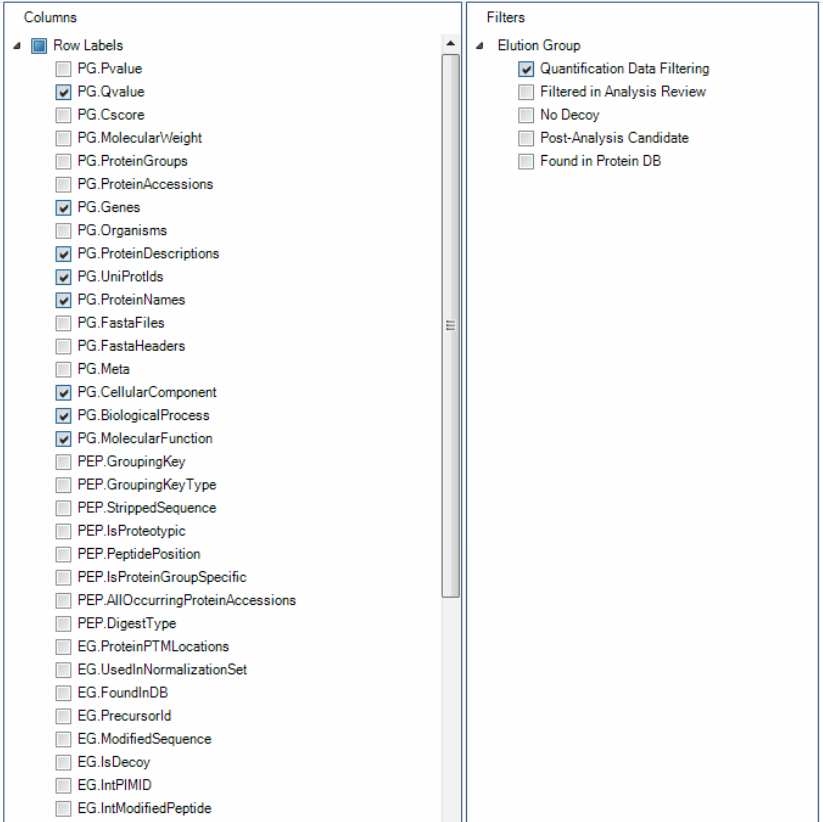

4. Select the following **Cell Values**:

- PG.NrOFPrecursorsIdentified
- PG.NrOFPrecursorsUsedForQuantification
- PG.Quantity.

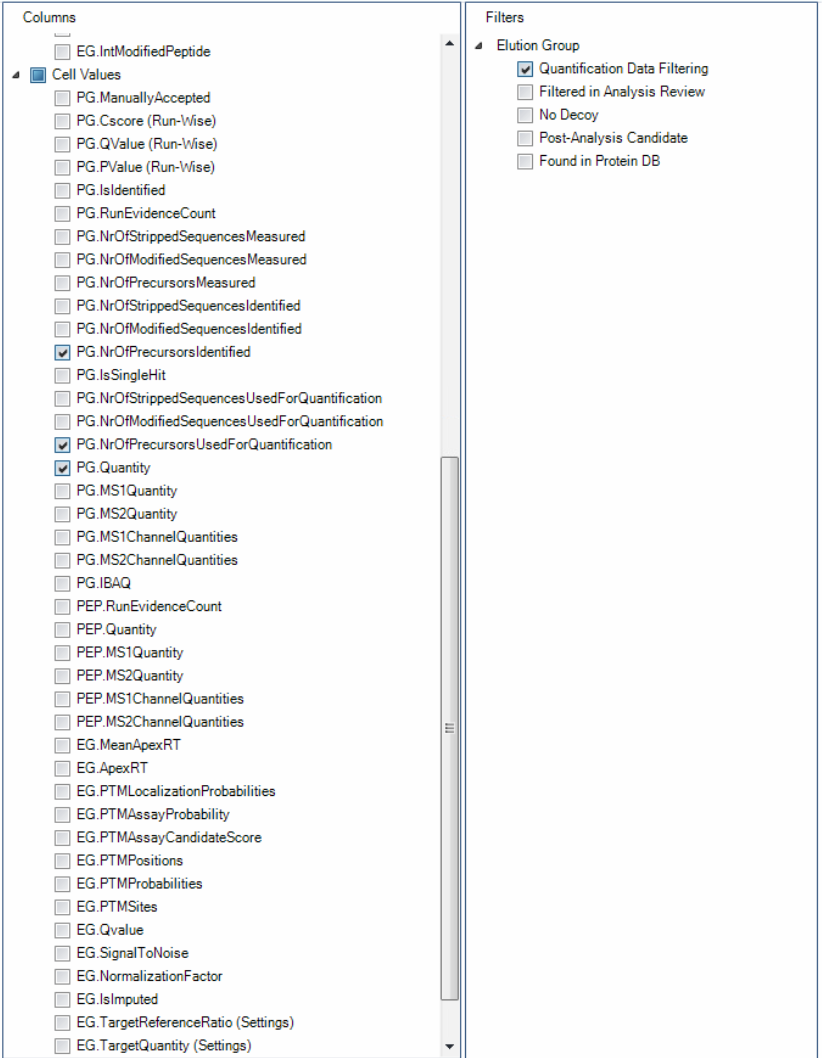

5. Select the following **Filters**:

- Quantification Data Filtering.

6. Select **Save As** and save the report scheme as “Protein_Quant_Pivot”.

#### **Chapter 9.2. Peptide Quantification Report**

1. Navigate to the **Report** tab to set up a custom report.

2. Select the **BGS Factory Report** under the **Run Pivot Report** dropdown to create a custom Peptide Quantification Pivot Report, “Peptide_Quant_Pivot”.

3. Select the following **Row Labels**:

- PG.Qvalue
- PG.Genes
- PG.ProteinDescriptions
- PG.UniProtIDs
- PG.ProteinNames
- PG.CellularComponent
- PG.BiologicalProcess
- PG.MolecularFunction
- PEP.PeptidePosition
- EG.PrecursorId
- EG.ModifiedSequence.

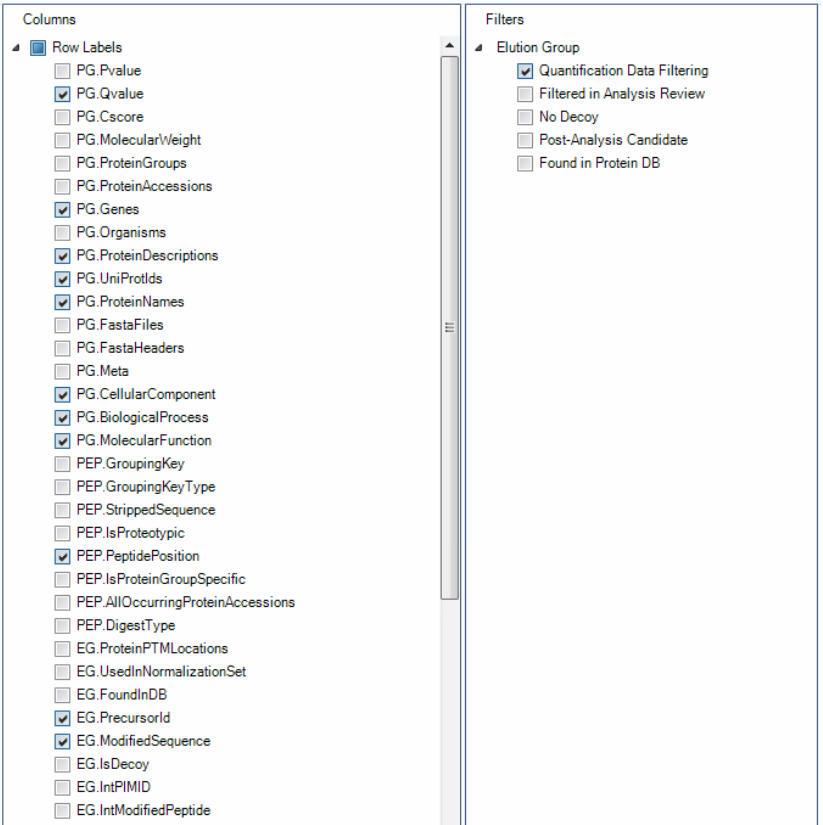

4. Select the following **Cell Values**:

- PEP.MS2Quantity
- EG.PTMProbabilities
- EG.PTMSites
- EG.TotalQuantity (Settings).

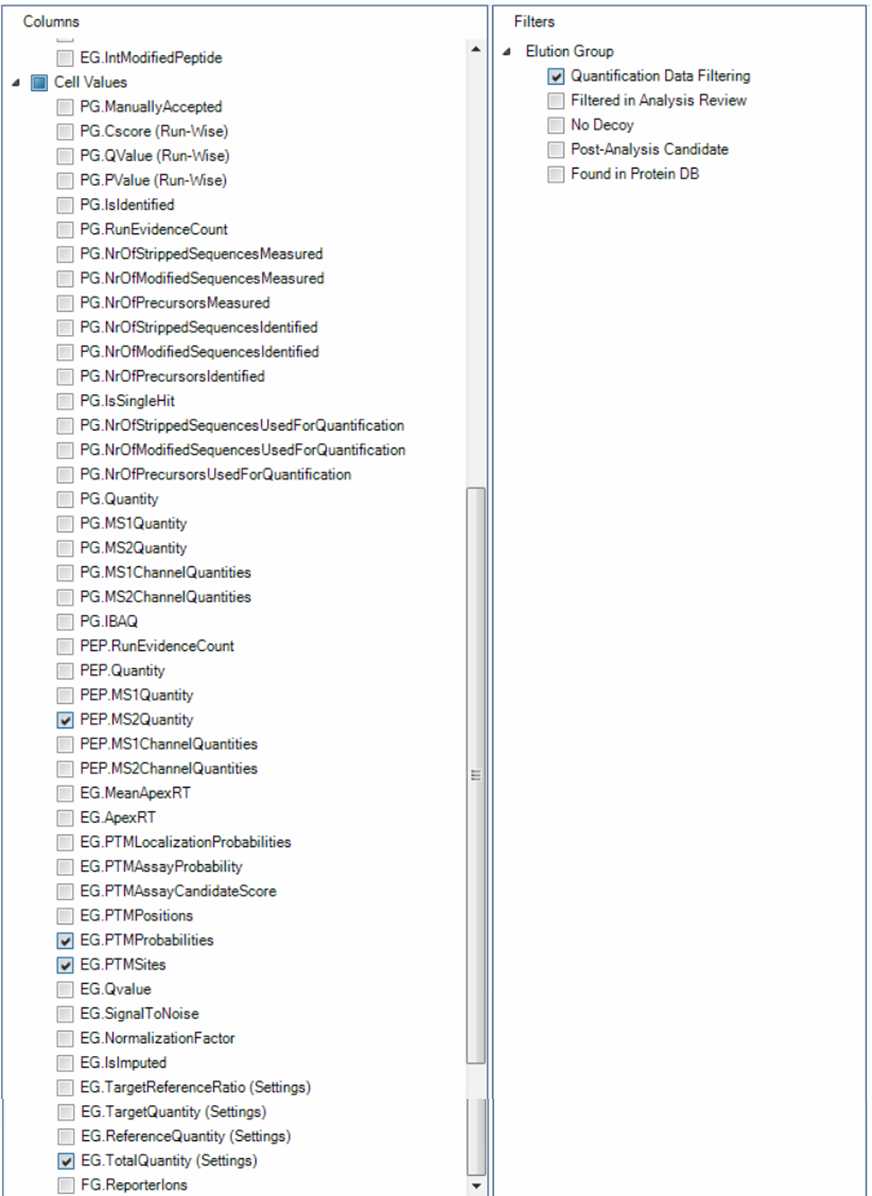

5. Select the following **Filters**:

- Quantification Data Filtering

6. Select **Save As** and save the report scheme as “Peptide_Quant_Pivot”.
